# TMT-Based Quantitative Proteomic Analysis of Intestinal Organoids Infected by *Listeria monocytogenes* with Different Virulence

**DOI:** 10.1101/2020.06.21.164061

**Authors:** Jie Huang, Cong Zhou, Guanghong Zhou, Keping Ye

## Abstract

*Listeria monocytogenes (Lm)* is an opportunistic food-borne pathogen that cause listeriosis. *L. monocytogenes* belonged to different serovars presents with different virulence in the host and caused different host reactions. To investigate the remodeling of host proteome by differently toxic strains, the cellular protein responses of intestinal organoids were analyzed using TMT labeling and high performance liquid chromatography-mass spectrometry. Quantitative proteomic analysis revealed 6564 differentially expressed proteins, of which 5591 proteins were quantified. The fold-change cutoff was set at 1.3 (*Lm* vs control), the virulent strain caused 102 up-regulated proteins and 52 down-regulated proteins, while the low virulent strain caused 188 up-regulated proteins and 25 down-regulated proteins. These identified proteins were involved in the regulation of essential processes such as biological metabolism, energy metabolism, and immune system process. Some selected proteins were screened by Real-time PCR and Western blotting. These results revealed that differently toxic *L. monocytogenes* induced similar biological functions and immune responses while had different regulation on differential proteins in the pathway.

## Introduction

*Listeria monocytogenes* is a gram-positive foodborne pathogen, which can lead to listeriosis ^[1]^. The infection causes a spectrum of illness, ranging from febrile gastroenteritis to invasive disease ^[2]^. Listeriosis is a relatively rare foodborne disease, while has a high mortality rate ranging from 20% to 30% ^[3]^. For example, the incidence of listeriosis was estimated to be 0.3 per 100000 persons in the United States, and the mortality was 21% in 2019 ^[4]^. Therefore, it is the second most frequent cause of foodborne infection-related deaths in Europe and USA ^[5]^. Moreover, listeriosis is mainly caused by contaminated food. In 2018, the listeriosis outbreak in South Africa was caused by ready-to-eat processed meat products, which resulted in 1034 cases of illness and 204 deaths ^[6, 7]^. Despite many achievements in food safety and laboratory diagnostics methods in developed countries, *L. monocytogenes* remains a major challenge in food industries and public health.

*L. monocytogenes* has 4 evolutionary lineages (I, II, III, and IV) based on multigene phylogenetic analyses, and was divided into 13 serotypes according to somatic O antigen ^[8]^. Of the 13 serotypes, over 98% of isolates from human listeriosis belong to serotypes within lineages I and II (1/2a, 1/2c, 1/2b, and 4b) ^[9]^. Food or food production environment was commonly contaminated with serotypes 1/2a and 1/2b, and clinical cases were mainly caused by virulent strain 4b, while low virulent strains were usually not pathogenic or weakly pathogenic, as serovar 4a ^[10]^. Besides, strains belonged to different serovars presents with different levels of virulence in the host and causes different host reactions, which due to their differences in growth and movement characteristics or expression of virulence factors ^[11, 12]^. In addition, *L. monocytogenes* will lose or weaken its toxicity due to the deletion of some virulence genes ^[13]^. There were many studies on the comparison of different toxic *L. monocytogenes*, mainly focusing on the following aspects: the genetic relationship between virulent and low-virulence strains ^[14]^, survival ability under stressful environments ^[15, 16]^, biological characteristics and pathogenicity in cell and mouse models ^[17, 18]^, expression of important virulence genes ^[19, 20]^, and host immune response ^[21]^, etc. Studies have shown that virulent strains are generally more pathogenic and invasive than attenuated strains. The *L. monocytogenes 10403s* in the 1/2a serotype strain can colonize the spleen and liver of mice in large quantities, and well invade Caco2 cells and replicate intracellularly ^[22]^. Moreover, strains of serotype 4a could cause infections, but could not establish long-term infections in macrophages ^[23]^. Compared with the 1/2a serotype strains, the 4a non-toxic strains had a shorter propagation time in the cells, but they all express similar metabolic related proteins.

*L. monocytogenes* is facultative intracellular pathogen that causes systemic infection by first invading the intestinal mucosal barrier ^[24]^. In the stage of intestinal infection, *L. monocytogenes* invaded intestinal epithelium by *L. monocytogenes* invasion protein InlA and zipper mechanism, entering Peyer’s patch through M cells, or leading the mislocalization of junctional proteins by Listeria adhesion protein ^[25, 26]^. The infection affected the normal intestinal renewal, causing the increased crypt depth and excessive proliferation, which damages the activity of intestinal stem cells ^[27]^.

Moreover, *L. monocytogenes* can immediately activate the host’s innate immune response when interacting with intestinal epithelial cells. Studies demonstrated that *L. monocytogenes* induced a dramatic innate inflammatory response in intestinal epithelial cells of germ-free, human E-cadherin transgenic mice ^[28]^. The innate immune activation was induced by pattern recognition receptors (PRRs) on epithelial cells, such as Toll-like receptors (TLRs) and nucleotide-binding oligomerization domain (NOD)-like receptors (NLRs). Many studies demonstrated the importance of TLR-mediated signaling in innate immune defense; however, the expression of TLR in the intestine was highly regulated and restricted to prevent exaggerated adaptive immunity to the intestinal microbiota ^[29, 30]^. Besides, several studies implicated that cytosolic proteins of NLRs contributed to the defense against intestinal bacterial infection. For example, it was found that mice lacking Nod2 have greater susceptibility to intestinal infection with *L. monocytogenes* ^[31]^. On the other hand, *L. monocytogenes* can employ strategies to evade or modulate immune defences. For example, *L. monocytogenes* modified bacterial ligands to avoid detection, modulated host signalling pathways to alter host innate defences ^[32]^. The detailed understanding of *L. monocytogenes*-host interactions and infection and immunity is essential for elucidating these mechanisms. Therefore, investigating the changes in the overall protein abundance of host cells by *L. monocytogenes* can help to understand the relationship between the pathogenicity and toxicity of the bacteria.

The majority of studies on *L. monocytogenes* infection have been conducted in animals such as mice, and single cell line models such as colon adenocarcinoma cells or macrophage cells ^[25, 33]^. Most knowledge of the innate and adaptive immune responses has been learned from experimental *L. monocytogenes* infections of mice ^[34]^. In addition to immune cells, intestinal epithelial cells were also involved in the defense against *L. monocytogenes* infections. Studies explored that host responses caused by *L. monocytogenes* included innate immune responses first activated by PRRs of intestinal epithelial cells, immune cells recruited by downstream cytokines and adaptive immune responses subsequently stimulated ^[32, 34]^. However, a single immune model or intestinal cell model both cannot completely reflect the damage of *L. monocytogenes* to the entire intestinal epithelium. Recently, intestinal organoid was emerging as a more effective infection model that reproduced the differentiation of intestinal epithelial cells, and showed the greatest similarity to the intestinal epithelium with respect to cell composition and structure ^[35]^. It was verified that organoids had been used to study the interaction between pathogens and host cells at the intestinal interface ^[36]^. Therefore, intestinal organoid is a suitable *L. monocytogenes* infection model for exploring the host response of non-immune cells.

Proteomics can provide protein information related to the biological metabolism and infection mechanism of the host or microorganism, which may be useful to further understand the interaction between the pathogenic microorganism and host ^[37]^. Recently, tandem mass tags (TMT)-based proteomic platforms have been used as one of the most robust proteomics techniques due to high sensitivity ^[38]^. It was shown that genomic and proteomics techniques were widely used to analyze the differences of strains in transcription and protein expression ^[39-41]^. Studies have shown that *L. monocytogenes* infection may have a major impact on host transcription and translation, cytoskeleton and connections, mitochondrial fission, host immune response, and apoptosis pathway ^[42, 43]^. However, researches on intestinal epithelial host responses caused by *L. monocytogenes* infection excluding the effects of immune cells is not yet comprehensive. Therefore, information on proteome changes in the infected intestinal organoids is necessary to understand host response of non-immune cells.

Here, intestinal organoids and two strains of *L. monocytogenes* (serotype 1/2a and 4a) were used, and the significant changes on global protein expression of infected intestinal organoids was described by a highly sensitive quantitative approach, including tandem mass tag (TMT) labeling and an LC-MS/MS platform combined with advanced bioinformatics analysis. Data revealed that major different proteins were involved in metabolic process, transcription and translation, and defense mechanisms (response to stimulus and immune system process). Furthermore, some significantly differentially expressed proteins (DEPs) related to host defence were further analyzed and validated.

## Materials and Methods

### Bacterial strains, Animals and Intestinal Organoids

The *Listeria monocytogenes 10403s* and *M7* was a gift of Prof. Weihuan Fang (Zhejiang University) ^[44, 45]^. Cryopreservation liquid of bacteria was transferred and scribed on PALCAM agar, and the plates were incubated at 37 °C for 48 h. Each single colony was picked out to 5 mL BHI broths supplemented with 5 µg/mL erythromycin and cultured with agitation at 37 °C for 16 h. The final concentration of the BHI broth was assessed by the plate count method.

4 weeks old, specific-pathogen-free (SPF) C57BL/6 mice were purchased from the Animal Research Centre of Yangzhou University. All the animal studies were approved by the Institutional Animal Care and Use Committee (IACUC) of Nanjing Agricultural University, and the National Institutes of Health guidelines for the performance of animal experiments were followed.

Intestinal organoids were cultured from intestines (mostly jejunum and ileal) of 4-week-old C57BL/6 mice, as described in Hou et al. After cervical dislocation, small intestine was removed and dissected immediately. Subsequently, intestine was flushed out with phosphate-buffered saline (PBS), and was cut into small pieces. Next, tissues were rocked in DPBS containing 2 mM EDTA for 30 min at 4 °C. After incubating, crypts were released by vigorously shaken, and cells were filtered through a 70-μm sterile cell strainer. Then, crypts were collected by centrifugation at 700 rpm for 5 min, mixed with Matrigel (Corning, USA) and then seed into a 24-well tissue culture plate. The plate was incubated for at least 15 min at 37 °C to polymerize. Finally, 500 mL of complete crypt culture medium was added to each well, which contained Advanced DMEM/F12 supplemented with penicillin–streptomycin, 10 mM HEPES, 2 mM glutamine, N2, B27 (Gibco, California, USA; Life Technologies, Carlsbad, California, USA), EGF (50 ng/mL, Peprotech, USA), R-spondin1 (500 ng/mL, Peprotech), Noggin (100 ng/mL, Peprotech), and Y-27632 (10 mM, Sigma, Germany). The medium was changed every 2–3 days, and organoids were passaged every 3–5 days.

### Experimental Design and *L. monocytogenes* Infection

Thirty four-week-old mice were randomly divided into 3 groups, 10 mice in each group, and placed in different cages. One group was inoculated with 1 × 10^9^ CFU of the virulent strain *L. monocytogenes 10403s* by oral gavage, the other group was also intragastrically inoculated with 1 × 10^9^ CFU of the attenuated strain *L. monocytogenes M7*, and another group was mockinfected with sterile PBS in the same manner. At 24, 72, and 96 hs after infection, mice were sacrificed, and CFUs in the intestine, liver and spleen were determined by dilution coating on Brain Heart Infusion plates with 5-µg ml–1 erythromycin and PALCAM agar plates. The body weight and survival rate were recorded for a week after inoculation.

Organoids were cultured in complete crypt culture medium at 37 °C in a 5% CO_2_-air atmosphere. To mimic the invasion of the intestine from the correct side, organoids were mechanical dissociated prior to infection, part of buds were fell off and thus exposed the lumen. After 3 days of passage, organoids were large enough to perform mechanical dissociation. Organoids were divided into 3 groups: one group was infected with 1 × 10^7^ CFU of the virulent strain *L. monocytogenes 10403s*; the other group was infected with 1 × 10^7^ CFU of the attenuated strain *L. monocytogenes M7*; and another group was mockinfected with sterile culture medium in the same mechanical dissociation. These were done by gently pipetting up and down with 10 mL pipettes to make a suitable wound ^[46, 47]^.

The detailed methods of organoids infection are listed as follows. First, *L. monocytogenes* was grown as described above and pelleted at 5000 rpm for 5 min. Subsequently, they were re-suspended in complete crypt culture medium to 10^7^ CFU/mL. Further, organoids were separated from Matrigel, and then re-suspended in complete crypt culture medium containing bacteria or not, and shook every 15 min at 37 °C. Specifically, infected organoids were incubated in complete culture medium with the indicated L. monocytogenes strain for 1 h while control organoids were only incubated with culture medium. After infection, organoids were centrifuged at 900 rpm for 5 min, and extracellular bacteria were removed by washing twice with DPBS. Finally, organoids were embedded into fresh Matrigel and cultured for 18 h, and the media was refreshed with penicillin-streptomycin media for the experiment.

### Protein Sample Preparation

Organoids were infected in three groups (virulent strain infection, attenuated strain infection and control) at two time points (1h after incubation and 18h after culture), a total of six groups; and three-well organoids were used as a sample, with three replicate samples in each group. Samples were sonicated three times on ice using a high intensity ultrasonic processor (Scientz) in lysis buffer (8 M urea, 1% Protease Inhibitor Cocktail). The remaining debris was removed by centrifugation at 12,000 g for 10 min. Finally, the supernatant was collected and the protein concentration was determined with BCA kit according to the manufacturer’s instructions. The aliquots were stored at −80 °C for further proteomic and Western blotting studies. The pooling of individual samples is a cost-effective approach for proteomic studies; therefore, four samples with equal amounts of protein from infected organoids or controls were mixed to obtain three virulent strain infection, three attenuated strain infection and three control samples.

### Trypsin Digestion

For trypsin digestion, the protein solution was reduced with 5 mM dithiothreitol for 30 min at 56 °C, and alkylated with 11 mM iodoacetamide for 15 min at room temperature in darkness. The protein sample was then diluted by adding 100 mM TEAB to urea concentration less than 2M. Finally, trypsin was added at 1:50 trypsin-to-protein mass ratio for the first digestion overnight and 1:100 trypsin-to-protein mass ratio for a second 4 h-digestion. Approximately 100 μg of protein for each sample was digested with trypsin for the following experiments

### TMT Labeling

After trypsin digestion, peptide was desalted by Strata X C18 SPE column (Phenomenex) and vacuum-dried. Peptide was reconstituted in 0.5 M TEAB and processed according to the manufacturer’s protocol for TMT kit. Briefly, one unit of TMT reagent (defined as the amount of reagent required to label 100 μg of protein) were thawed and reconstituted in acetonitrile. The peptide mixtures were then incubated for 2 h at room temperature and pooled, desalted and dried by vacuum centrifugation. Prepared samples were stored at –80 °C until liquid chromatography-mass spectrometry (LC–MS/MS) analysis.

### HPLC Fractionation

The samples were fragmented into a series of fractions by high pH reverse-phase HPLC using Agilent 300Extend C18 column (5 μm particles, 4.6 mm ID, and 250 mm length). Briefly, peptides were first separated with a gradient of 8% to 32% acetonitrile (pH 9.0) over 60 min into 60 fractions. Then, the peptides were combined into 18 fractions and dried by vacuum centrifuging.

### LC-MS/MS

The peptides were dissolved in liquid chromatography mobile phase A (0.1% formic acid and 2% acetonitrile) and separated using the EASY-nLC 1000 ultra-high performance liquid system. Mobile phase B is an aqueous solution containing 0.1% formic acid and 90% acetonitrile. Liquid phase gradient setting as follows: 0 ∼ 42 min, 6% −22% B; 42 ∼ 54 min, 22% −30% B; 54 ∼ 57 min, 30% −80% B; 57 ∼ 60 min, 80% B, all at a constant flow rate of 500 nL / min.

The peptides were subjected into the NSI ion source for ionization, and then analyzed by Orbitrap Fusion Lumos™ mass spectrometry. The ion source voltage was set to 2.4 kV, the peptide precursor ions and their secondary fragments were detected and analyzed using high-resolution Orbitrap. The m / z scan range of full scan was 350 to 1550, and the resolution of the complete peptide detected in Orbitrap was 60,000; the scan range of the second-level mass spectrometer was fixed at 100 m/z, and the resolution was set to 15,000. The data acquisition mode used a data-dependent scanning (DDA) program. In order to improve the effective utilization of the mass spectrum, the automatic gain control (AGC) was set to 5E4; the signal threshold was set to 10000 ions/ s; the maximum injection time was set to 60 ms; and the dynamic exclusion time of the tandem mass spectrometry scan was set to 30 seconds to avoid precursor ion Repeat the scan.

### Data Analysis

The resulting MS/MS data were processed using Maxquant search engine (v.1.5.2.8). Tandem mass spectra were searched against SwissProt Mouse database concatenated with reverse decoy database. An anti-library was added to calculate the false positive rate (FDR) caused by random matching, and a common contamination library was added to the database to eliminate the impact of contaminated proteins in the identification results. Trypsin/P was specified as cleavage enzyme allowing up to 2 missing cleavages. The minimum length of the peptide was set to 7 amino acid residues; the maximum number of modifications of the peptide was set to 5; the mass error of the primary precursor ion of First search and Main search was respectively set to 20 ppm and 5 ppm; and the mass error of the secondary fragment ion was 0.02 Da.

The cysteine alkylation was set as a fixed modification, and the variable modification was the oxidation of methionine, acetylation at the N-terminus of the protein, and deamidation (NQ). The quantitative method was set to TMT-6plex, FDR for protein identification and PSM identification was set to 1%, and minimum score for peptides was set > 40.

### Real-Time RT –PCR

Total RNA was respectively extracted from organoid sample using RNAiso Plus (Takara, Beijing, China). Following, reverse transcription of the RNA was performed. With the primers listed in Table 1, master mix was used to yield a final volume of 20 µL (Takara). The thermal cycling conditions were 5 min at 95 °C, followed by 40 cycles of 15 s at 95 °C and 34 s at 60 °C using an Applied Biosystems 7500 real-time PCR system as described previously (Hou et al., 2018). The mRNA expression level of each target gene was normalized to the expression level of GAPDH, the expression levels of uninfected organoids comparing the expression levels of infected organoids were normalize as 1 which was analyzed by ΔΔCt. All real-time PCR reactions were performed in triplicate. Primer sequence of target and reference genes were shown as Table 1.

**Table 1.**
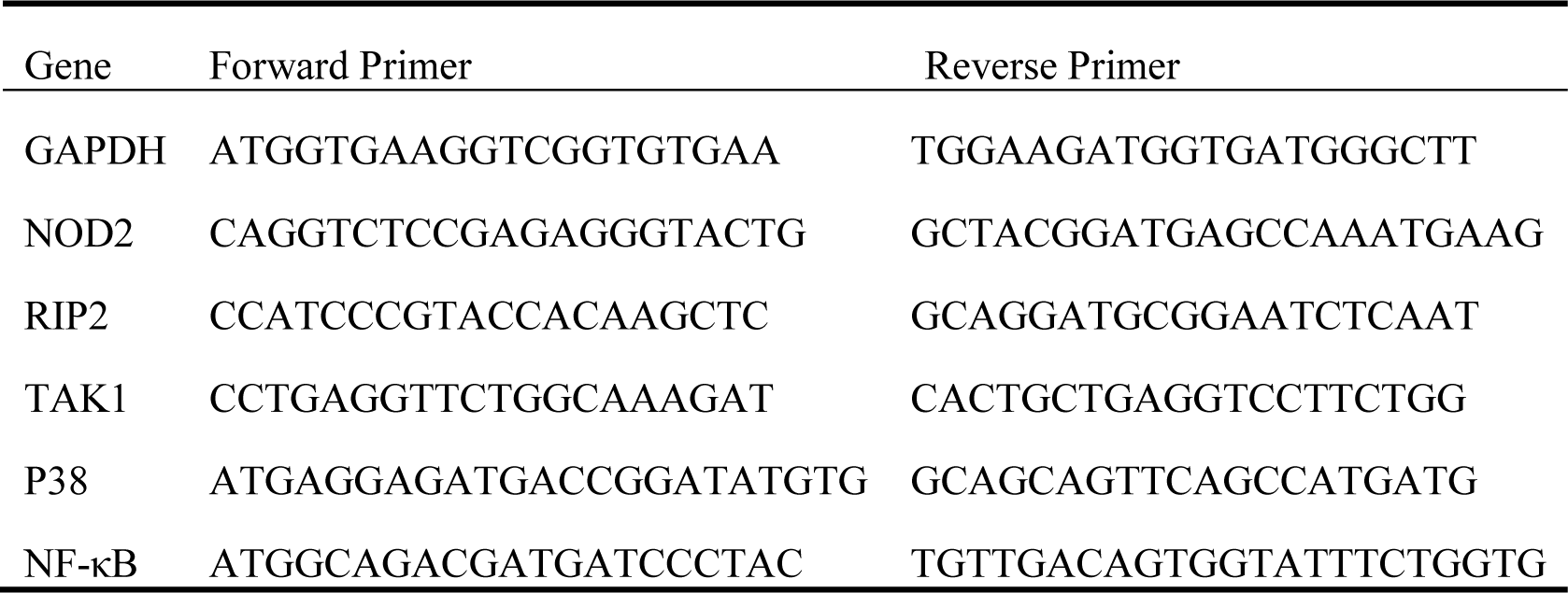
Primer sequence of target and reference genes

### Western Blot Analysis

Organoids were lysed in RIPA buffer (50mM Tris-HCl, pH 7.4, 1% NP-40, 150mM NaCl) containing a protease inhibitor cocktail (Thermo FisherScientific). Protein concentrations were detected using a BCA protein quantification kit (Thermo Fisher Scientific). Equal amounts of protein were separated by 10% SDS-PAGE, and then transferred to PVDF membranes (Millipore, China). After blocking with 5% non-fat milk in TBS containing 0.1% Tween-20, the membranes were probed with the appropriate antibody. The following antibodies were used for Western blot analysis as follows: rabbit anti-NOD2 (1:400), anti-GAPDH (1:1,000, SAB4300645, Sigma). After washing, the membranes were incubated with goat anti-rabbit secondary antibodies (1:10000). Finally, Blots were developed using efficient chemiluminescence (ECL) kit and light emission was captured using the Versa DOC 4000 imaging system. Use Quantity one software to analyze the gray value of the band.

## Results

### *Listeria monocytogenes* infection and clinical signs

After 3 days of acclimation, the mice were in good condition with normal activity. After randomized intragastric administration, the control group was in a normal state, and both the group of virulent strain *L. monocytogenes 10403s* and low virulent strain *L. monocytogenes M7* showed abnormal state and deaths with different degrees. As shown in Figure 1A, within 7 days after gavage, the overall weight of the control group showed an upward trend, with the largest increase in the 4th to 5th days, and there was no death during the entire process. The weight of the virulent group showed a trend of decreasing first and then increasing. It continued to decrease in the first 4 days and increased after the 5th day. Compared with the control group, the mice died on the first day after gavage, and the survival rate continued to decline, and did not change after the 5th day.

**Figure 1.**
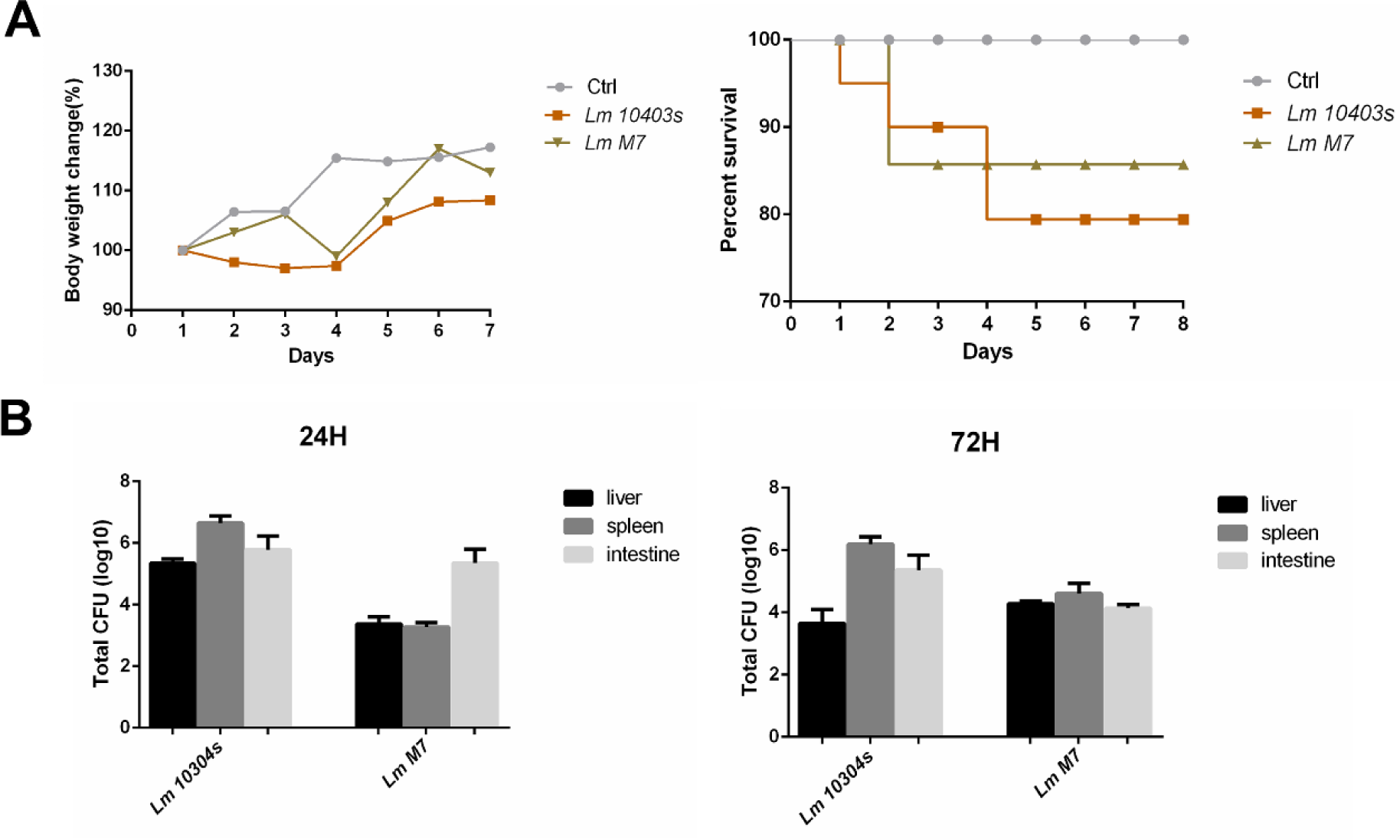
*Listeria monocytogenes* infection in mice. A. The body weight change rate and survival rate of mice 7 days after gavage. B. Bacterial load after 24h and 72h infection

The overall weight of the low virulent strain group fluctuated, and the change was large, which decreased on the 3rd day and increased on the 4th-6th days; the group only died on the 2nd day, and the final survival rate was lower than the control group but higher than the virulent group. 24 h after infection, two strains of *L. monocytogenes* can be respectively detected in the intestine, liver, and spleen (Figure 1B). The main lesions of infected mice comprised partial hemorrhage in small intestine, longer colon length and enlarged spleen, and lesions caused by the virulent strain were more serious.

The results showed that mice in the infected groups were successfully infected with two stains of *L. monocytogenes*, and they both induced short-term body weight changes and varying degrees of death. In contrast, highly toxic *L. monocytogenes 10403s* resulted in more serious infections and lower survival rates.

### Quality validation of the proteomic data

To reveal the changes in the protein levels under *L. monocytogenes* infections with different toxicity, an integrated proteomic approach was performed using organoids (control and two infected group). The statistical information of the treatment groups and differential proteins were shown in Table 2 and Table 3. As shown in Figure 2A, the Pearson coefficient between all replicate samples was greater than 0.6, which met the biological repeat quantitative consistency standard; it indicated that the correlation coefficient of 18 experimental samples displayed good repeatabilities. Two important parameters, including peptide length and peptide mass, were analyzed to verify the quality of mass spectrometry data. The data showed that the sample preparation met the standard requirements, most of the peptides were distributed between 7-20 amino acids, which is in accordance with the general law based on trypsin digestion and HCD fragmentation, and the first-order mass error of most spectra is within 10 ppm (Figure 2B and C).

**Table 2.** Treatment conditions of 6 treatment groups

| Groups | 1 h | 18 h |
| --- | --- | --- |
| Control | A | B |
| *L. monocytogenes 10403s* | C | D |
| *L. monocytogenes M7* | E | F |

**Table 3.** Summary of Differentially expressed proteins

| Comparison group | up-regulated (>1.3) | down-regulated (<1/1.3) |
| --- | --- | --- |
| B/A | 103 | 146 |
| C/A | 37 | 2 |
| E/A | 6 | 12 |
| D/B | 102 | 52 |
| F/B | 188 | 25 |
| C/D | 486 | 388 |
| C/E | 107 | 12 |
| D/F | 19 | 121 |
| E/F | 386 | 595 |

**Figure 2.**
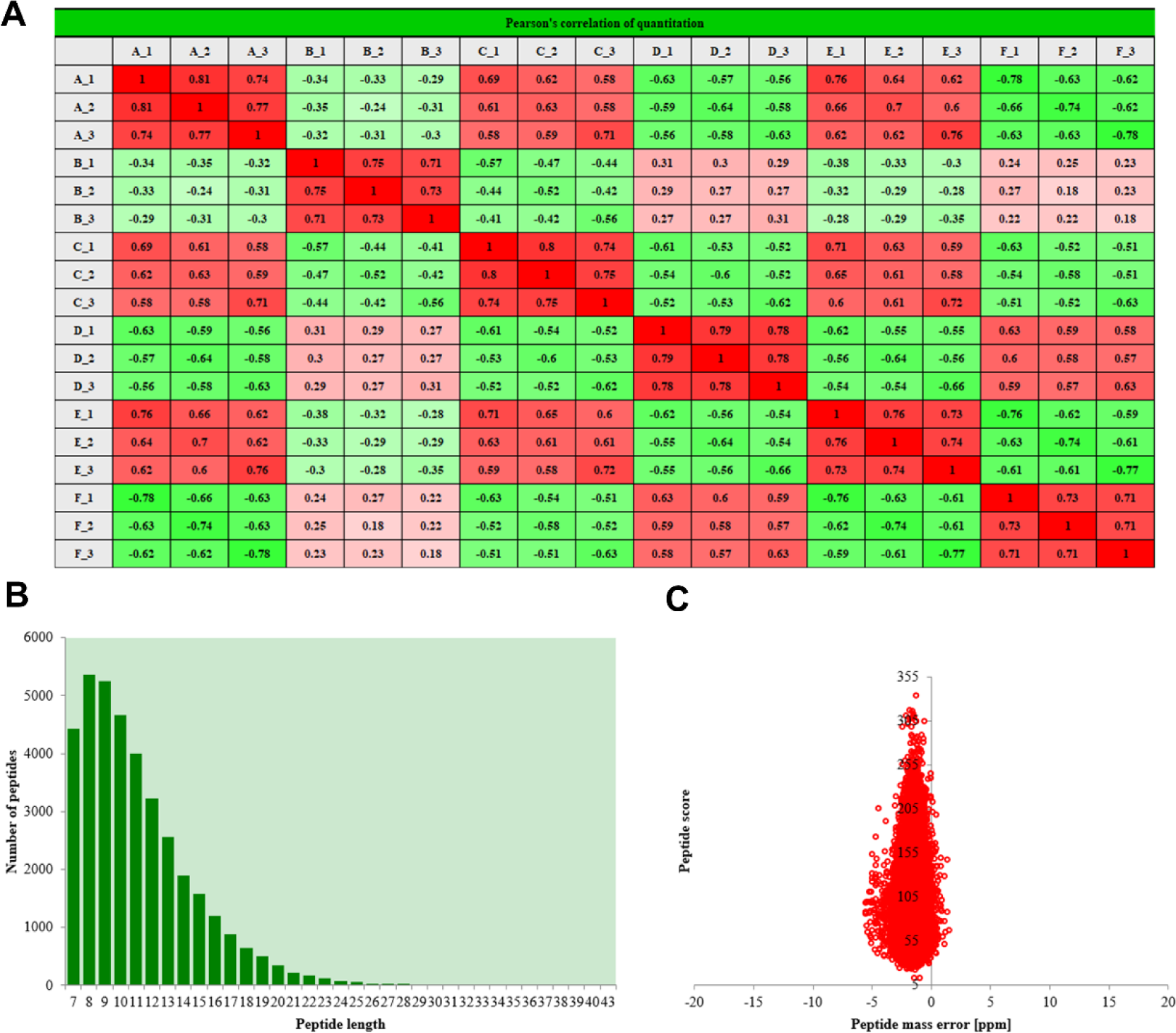
Experimental strategy for quantitative proteome analysis and QC validation. A. Pearson’s correlation of protein quantitation. B. Length distribution of all identified peptides. X-axis: No. of Peptide; Y-axis: Peptide length. C. Mass delta of all identified peptides. X-axis: Peptide Score; Y-axis: Peptides mass delta.

### Analysis of the DEPs under *L. monocytogenes 10403s* and *L. monocytogenes M7* infections

In this study, we performed a quantitative analysis of the overall proteome of small intestinal infected organoids. Altogether, 6564 proteins were identified, of which 5591 proteins were quantified. For the differentially expressed proteins (DEPs), the cutoff criteria considered was set with p value < 0.05 and the infection vs control group ratio >1.3-fold difference. Based on our previous experimental results, three groups of differential proteins cultured for 18 h after incubation were subsequently analyzed (B, D, F group). Among the quantitative proteins, 102 up-regulated and 52 down-regulated proteins were identified under *L. monocytogenes 10403s* infection, while 188 up-regulated and 25 down-regulated proteins were identified under *L. monocytogenes M7* infection. The differential expressed proteins between *L. monocytogenes 10403s* and *L. monocytogenes M7* infection were also calculated, resulting in 4 up-regulated and 58 down-regulated proteins (Figure 3). Table S1 (Supplementary Materials) presents relative expression of DEGs in *L. monocytogenes 10403s* vs Control (D/B) and *L. monocytogenes M7* vs Control (F/B). The protein ratio were expressed as *L. monocytogenes* infection vs control.

**Figure 3.**
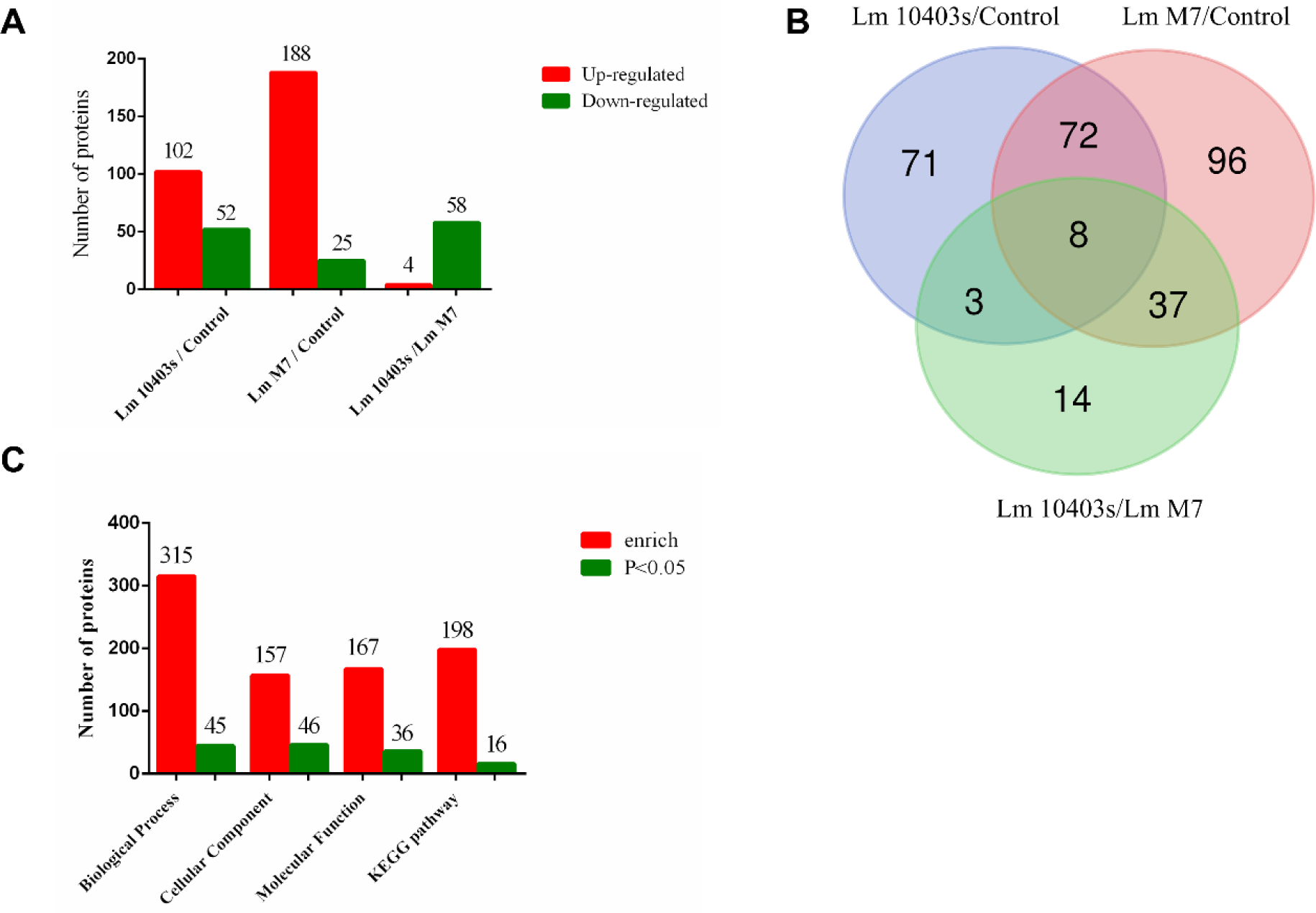
The numbers of DEPs in different comparisons. A. The numbers of the up-and down-regulated proteins in each comparison. B. Venn diagram of DEPs in different comparisons. C. Statistical overview of the bioinformatics of DEPs obtained in the combination of *L. monocytogenes 10403s* infection group and *L. monocytogenes M7* infection group.

### Classification of the DEPs under *L. monocytogenes 10403s* and *L. monocytogenes M7* infections

To further understand the DEPs in *L. monocytogenes*-infected intestinal organoids, functional classification was performed from the Gene Ontology (GO) and subcellular structure localization (Figure 4). GO was divided into three main categories: biological process, cellular component and molecular function, which can explain the biological role of proteins from different angles.

**Figure 4.**
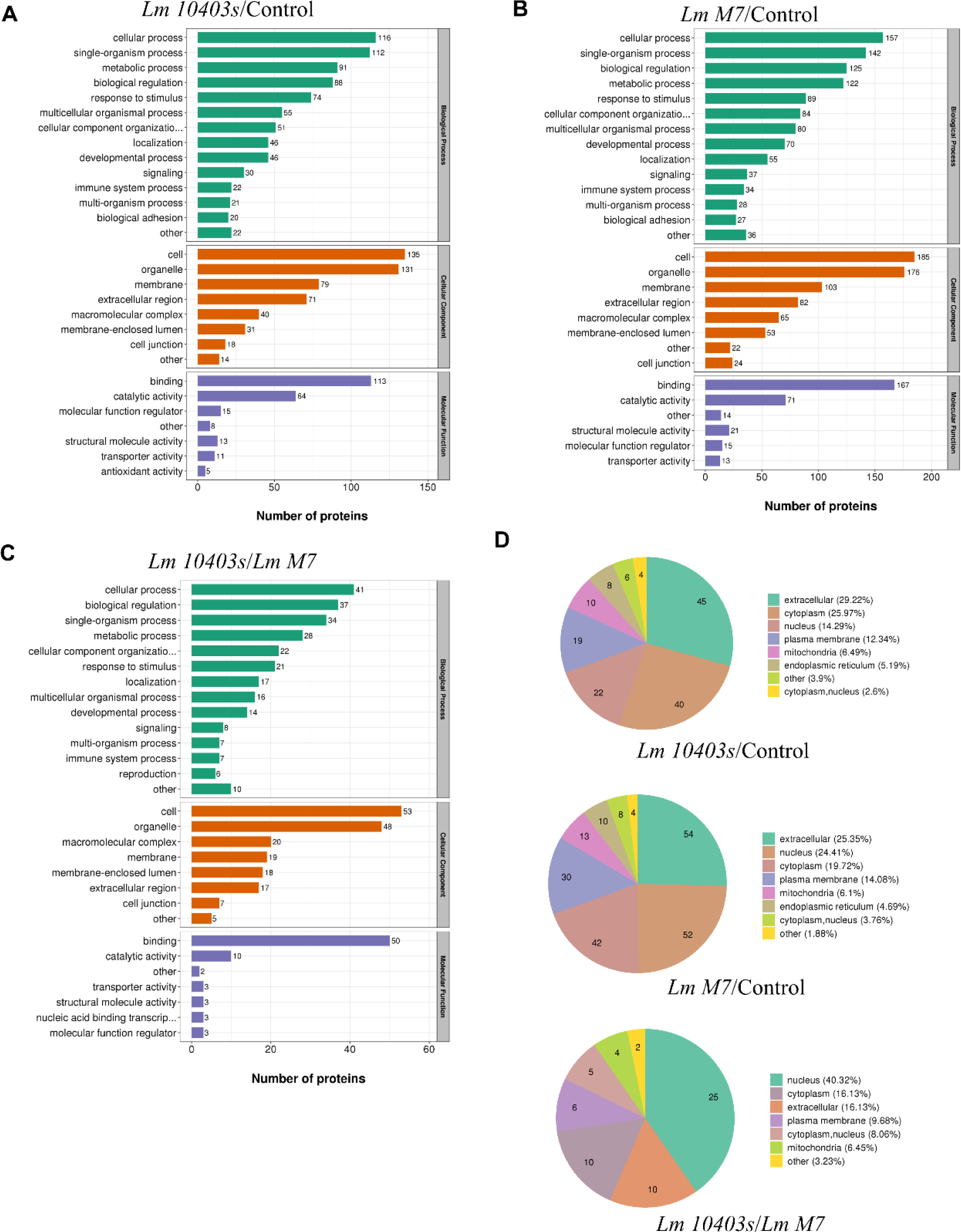
GO analysis and subcellular locations of DEPs in different comparisons. A. GO analysis of regulated DEPs in *L. monocytogenes 10403s* vs Control. B. GO analysis of regulated DEPs in *L. monocytogenes M7* vs Control. C. GO analysis of the up-regulated DEPs in *L. monocytogenes 10403s* vs *L. monocytogenes* M7. D. GO analysis of the down-regulated DEPs in *L. monocytogenes 10403s* vs *L. monocytogenes* M7. All proteins were classified by GO terms. X-axis: Number of DEPs. E. Subcellular locations of the DAPs in different comparisons.

Under the *L. monocytogenes 10403s* infection, the main biological processes of DEPs included single-organism process, biological regulation and metabolism, defense response from stimulation and immunity; cell composition mainly included membrane and macromolecular complex, and molecular function mainly included binding and catalytic activity (Figure 4A). Under *L. monocytogenes M7* infection, the main biological processes of DEPs were the same as those of *L. monocytogenes 10403s*. The main conclusions of cell composition and molecular function were also similar to those of *L. monocytogenes 10403s* (Figure 4B). In the comparison between *L. monocytogenes 10403s* and *L. monocytogenes M7*, up-regulated proteins were grouped into single–organism process, membrane and binding; while down-regulated proteins were related to biological regulation and metabolism, immune system process, macromolecular complex and catalytic activity (Figure 4C and D).

In the subcellular localization of DEPs, the host cell proteins infected by *L. monocytogenes 10403s* were mainly distributed in the extracellular and cytoplasm, and the proteins of *L. monocytogenes M7* infection were mainly distributed in the extracellular and nucleus. The DEPs of comparing two infections were mainly distributed in the nucleus (Figure 4E).

### Enrichment analysis of the DEPs under *L. monocytogenes 10403s* and *L. monocytogenes M7* infections

To find out whether DEPs had a significant enrichment trend in certain functional types, enrichment analysis of GO classification and protein domain were performed in each comparison group (Figure 5). Fisher’s exact test p-value (-log10 [p-value]) was used to evaluate the enrichment level of DEPs; the larger the p-value, the more differentially expressed proteins enriched in this category.

**Figure 5.**
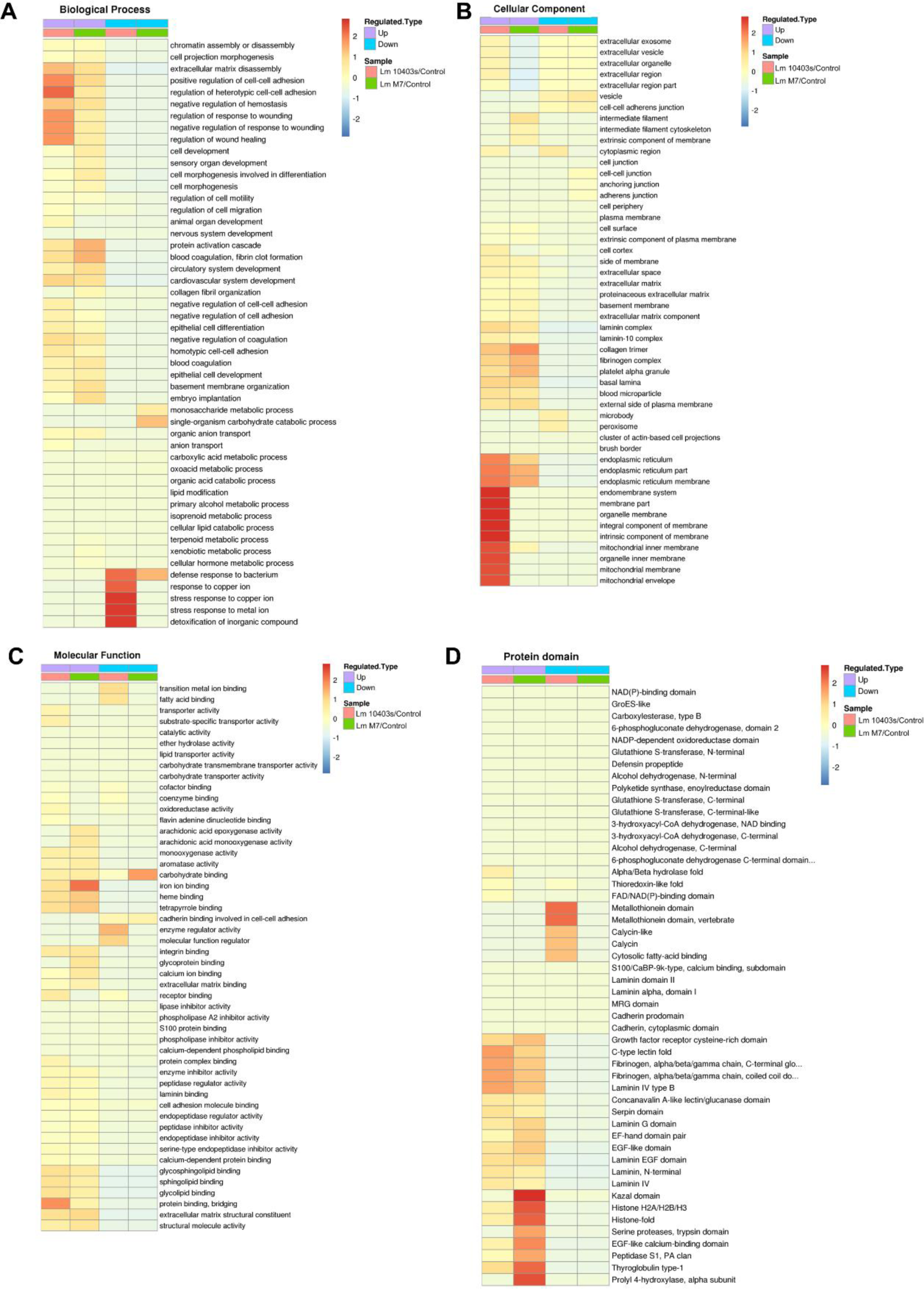
GO and protein domain enrichment analysis of the DEPs. Heatmaps showed the enrichments of the DEPs in different comparisons with GO annotation belonging to biological process (A), cellular component (B), and molecular function (C). D. Significantly enriched protein domains of the DEPs.

The first was the enrichment analysis of GO classification. Under *L. monocytogenes 10403s* infection, the significantly enriched biological process were mainly associated with defense response to bacterium, antimicrobial humoral response, extracellular matrix disassembly; the significantly enriched cellular component were related to extracellular matrix, basement membrane; the significantly enriched molecular function were mainly correlated with extracellular matrix structural constituent, oxidoreductase activity and various bindings, such as glycosphingolipid binding. Besides, the significantly enriched domain terms were Laminin EGF domain, Laminin and Fibrinogen, and Metallothionein domain.

Under *L. monocytogenes M7* infection, the significantly enriched biological process were mainly associated with protein activation cascade, antimicrobial humoral response, extracellular matrix disassembly; the significantly enriched cellular component were basically the same as *L. monocytogenes 10403s*; the significantly enriched molecular function were mainly correlated with extracellular matrix structural constituent, iron ion binding and structural molecule activity. Moreover, the top three significantly enriched domain terms were same as *L. monocytogenes 10403s*, while the remaining domains included Histone.

In the comparison between *L. monocytogenes 10403s* and *L. monocytogenes M7*, the significantly enriched biological processes were mainly associated with negative regulation of gene expression and biosynthetic process, cellular macromolecular complex assembly. The significantly enriched cellular components were related to nucleosome and DNA packaging complex. The significantly enriched molecular functions were mainly correlated with chromatin DNA binding and nucleosomal DNA binding. In addition, the significantly enriched domain term were Histone and Histone-fold.

### KEGG analysis of the DEPs under *L. monocytogenes 10403s* and *L. monocytogenes M7* infections

All DEPs in *L. monocytogenes 10403s* infection group and *L. monocytogenes M7* infection group were put together for KEGG analysis. Figure 6A showed all the KEGG pathways enriched. Combining with Figure 3C, 301 differential proteins participated in 198 KEGG pathways, of which only 16 were significantly enriched. Figure 6B was the percentage of these enriched KEGG pathways. The larger the proportion, the more proteins involved in this pathway. The most significant enrichment was chemical carcinogenesis, which reflected the genotoxic and non-genotoxic effects of *L. monocytogenes* on the host. The main significant enrichments were some metabolic related pathways, such as Retinol metabolism, steroid hormone biosynthesis, and drug metabolism-cytochrome P450. In addition, fat digestion and absorption, glycolysis/gluconeogenesis and other metabolic pathways were also been enriched. Besides, the DEPs reflected the ability of *L. monocytogenes* to adhere and damage during the invasion were significantly involved in the small cell lung cancer, Amoebiasis and some other host responses, such as the ECM-receptor interaction, Complement and coagulation cascades, and HIF-1 signaling pathway. Moreover, it could be seen from Figure 6B that some signaling pathways related to host defense and immune response were enriched, which didn’t contain many differential proteins, such as apoptosis and autophagy, Ferroptosis, and NOD-like receptor signaling pathway, etc. Figure 7 showed that KEGG pathway of *L. monocytogenes 10403s* and *L. monocytogenes M7* infection group were mainly related to invasion, host response and metabolism. Among the KEGG pathways to which up-regulated protein enriched in the *L. monocytogenes 10403s* infection group, the pathways related to bacterial adhesion and invasion included Amoebiasis, small cell lung cancer, chemical carcinogenesis and focal adhesion. In addition, it also included some host response, such as ECM-receptor interaction, complement and coagulation cascades and PI3K-Akt signaling pathway. Moreover, KEGG pathways related to metabolic processes contained steroid hormone biosynthesis, retinol metabolism and drug metabolism – cytochrome P450.

**Figure 6.**
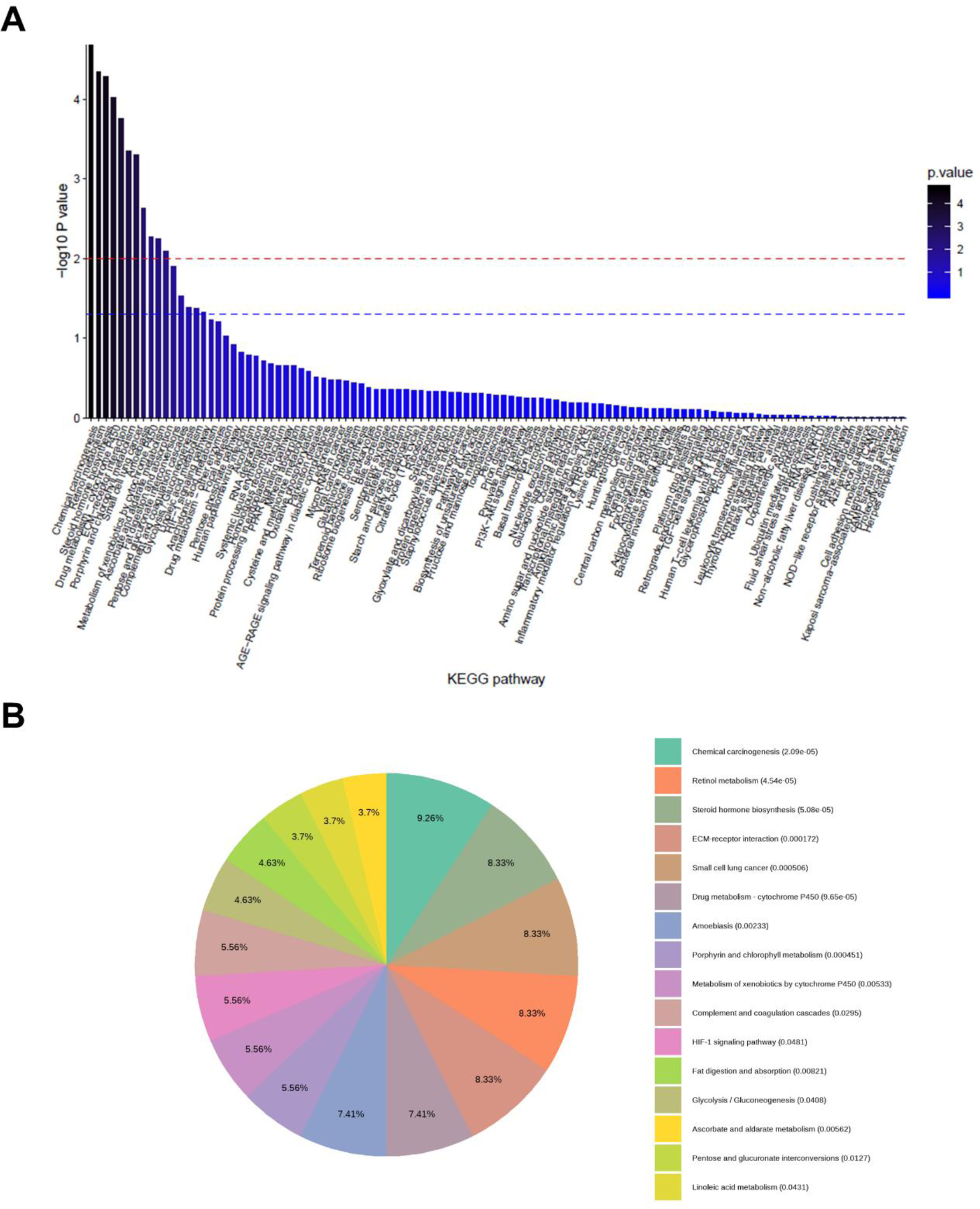
KEGG enrichment analysis of all DEPs obtained in the combination of *L. monocytogenes 10403s* infection group and *L. monocytogenes M7* infection group All differentially expressed proteins in *L. monocytogenes 10403s* infection group and *L. monocytogenes M7* infection group were put together for KEGG analysis. A. Columnar Section of KEGG enrichment analysis results. B. Pie chart of KEGG enrichment analysis results.

**Figure 7.**
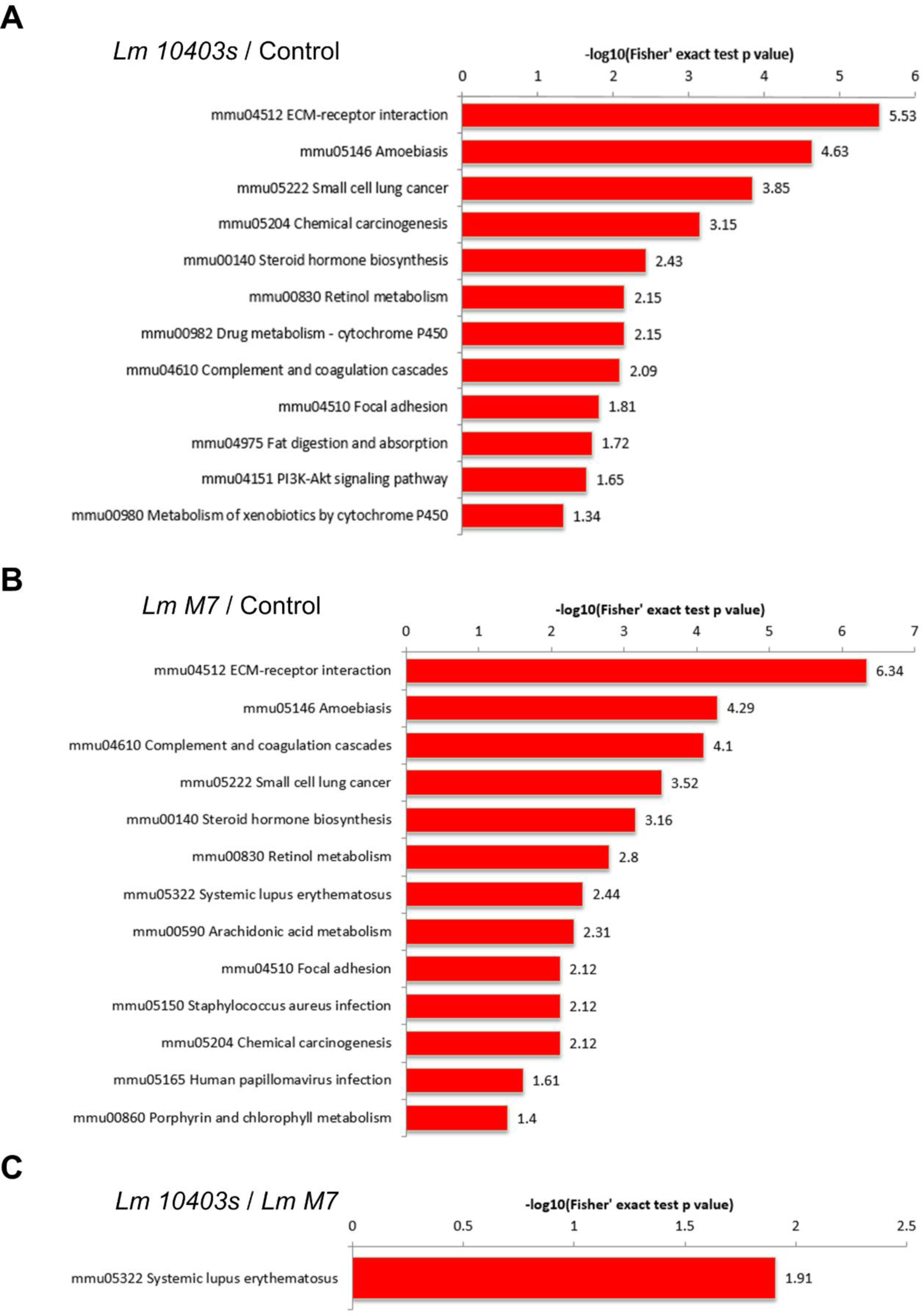
KEGG enrichment analysis of the DEPs. A. Significantly enriched KEGG terms of the DAPs in the *L. monocytogenes 10403s* vs control comparison. B. Significantly enriched KEGG terms of the DAPs in the *L. monocytogenes M7* vs control comparison. C. Significantly enriched KEGG terms of the DAPs in the *L. monocytogenes 10403s* vs *L. monocytogenes M7* comparison.

Similar results were found in the KEGG pathway enriched in the *L. monocytogenes M7* infection group. Up-regulated proteins were enriched in the same three pathways related to bacterial adhesion and invasion as *L. monocytogenes 10403s*, involved in ECM-receptor interaction and complement and coagulation cascades, and enriched in three host lipid metabolism-related metabolic processes. However, unlike the virulent strain *L. monocytogenes 10403s*, the upregulated protein of *L. monocytogenes M7* were contributed in systemic lupus erythematosus, and the downregulated proteins were only enriched in glycolysis/gluconeogenesis.

In our intestine organoid model, the innate immune response was the primary host defense response caused by *L. monocytogenes*. Therefore, we selected the DEPs of five pathways in the *L. monocytogenes 10403s* vs control group and *L. monocytogenes M7* vs control group, which were related to the immune system process. Then these proteins were accurately analyzed at 1.3-fold and 1.2-fold differential folds. The five pathways included ECM-receptor interaction, complement and coagulation cascade, HIF-1 signaling pathway, Ferroptosis and NOD-like receptor signaling pathway.

Among the ECM-receptor interactions, the significantly upregulated proteins which were mutual in the *L. monocytogenes 10403s* and *M7* groups were Fn1, Lamc1, Lama1, Lamb1, Col4a2, Col4a1, Lamb2, Hspg2, and Agrn. Besides, Agrn was up-regulated but not significant in the *L. monocytogenes 10403s* group, while 1.3-fold significantly up-regulated in the *L. monocytogenes M7* group, and 1.2-fold significant in the *L. monocytogenes 10403s* vs *M7* comparison group. It showed that *L. monocytogenes M7* could significantly increase Agrn 1.3-fold, and the degree of upregulation was1.2-fold more significant than *L. monocytogenes 10403s*.

For the complement and coagulation cascades, six differential proteins were both identified in the *L. monocytogenes 10403s* and *M7* infection groups, Plg, Fgb, Fga, Fgg, Clu, and C3. These differential proteins were significantly up-regulated in the *L. monocytogenes 10403s* and *M7* infection groups, while not significantly changed in the comparison group. In addition, F10 was significantly changed in the *L. monocytogenes 10403s* vs *L. monocytogenes M7* comparison group, which was significantly up-regulated 1.3-fold in the *L. monocytogenes M7* group, while was down-regulated but not significant in the *L. monocytogenes 10403s* group. This showed that effect of *L. monocytogenes M7* on F10 was opposite to *L. monocytogenes 10403s*, and the degree of activation was significantly higher than *L. monocytogenes 10403s*.

In the NOD-like receptor signaling pathway, the differential proteins shared by the *L. monocytogenes 10403s* and *M7* infection groups were significantly down-regulated. Among them, Nod2 and Pycard were 1.3-fold significantly reduced in both *L. monocytogenes 10403s* group and *L. monocytogenes M7* group, and Nampt was 1.2-fold significantly reduced in both group. In addition, the 1.2-fold significantly up-regulated proteins in the *L. monocytogenes 10403s* group were Vdac1, Vdac2, and Vdac3, which were also up-regulated in the *L. monocytogenes M7* group but the changes were not significant. This suggested that one of the differences between the effects of *L. monocytogenes 10403s* and *L. monocytogenes M7* on the NOD-like signaling pathway was the activation of these proteins. In particular, Txn2 and Nek7 were significantly down-regulated in the *L. monocytogenes 10403s* vs *L. monocytogenes M7* comparison group, they were down-regulated in the *L. monocytogenes 10403s* group and up-regulated in the *L. monocytogenes M7* group, which both were not significantly. It indicated that the activation of *L. monocytogenes M7* on these proteins was significantly different from *L. monocytogenes 10403s*.

In the HIF-1 signaling pathway, the significant differentially upregulated differential proteins shared by the *L. monocytogenes 10403s* and *M7* infection groups were Tfrc, Tf, Hkdc1 and Cdkn1b, and the significantly downregulated protein was Eno1. However, only Slc2a1 changed significantly in the *L. monocytogenes 10403s* vs *L. monocytogenes M7* comparison group, which was significantly increased.

For Ferroptosis, the significantly upregulated proteins shared by *L. monocytogenes 10403s* and *M7* infection groups were Acsl5, Tf, Acsl1, Tfrc, and Lpcat3; and the significantly down-regulated proteins were Fth1 and Ftl1, which both did not change significantly in the comparison group. In addition, *L. monocytogenes 10403s* 1.2-fold significantly increased Gpx4, Vdac3 and Vdac2, which were up-regulated but not significant in the *L. monocytogenes M7* group.

### Confirmation of proteomic data by RT-PCR and western blot analysis

Among the five related pathways, only the differential proteins in the NOD-like receptor-signaling pathway were down-regulated. This showed that the effect of *L. monocytogenes* infection on the NOD-like receptor-signaling pathway is mainly to inhibit or reduce the differential proteins. Among them, Nod2 is expressed in Paneth cells and stem cells in the intestine, and very important for intestinal immunity. Therefore, the gene and protein level of the Nod2 pathway protein were verified. The results were shown in Figure 8. *L. monocytogenes* with different toxicity significantly increased the mRNA expression level of Nod2, while it was down-regulated at the protein level, consistent with the proteome results. However, for other proteins in the Nod2 pathway, *L. monocytogenes* 10403s significantly down-regulated RIP2, TAK1, P38 and NF-κB.

**Figure 8.**
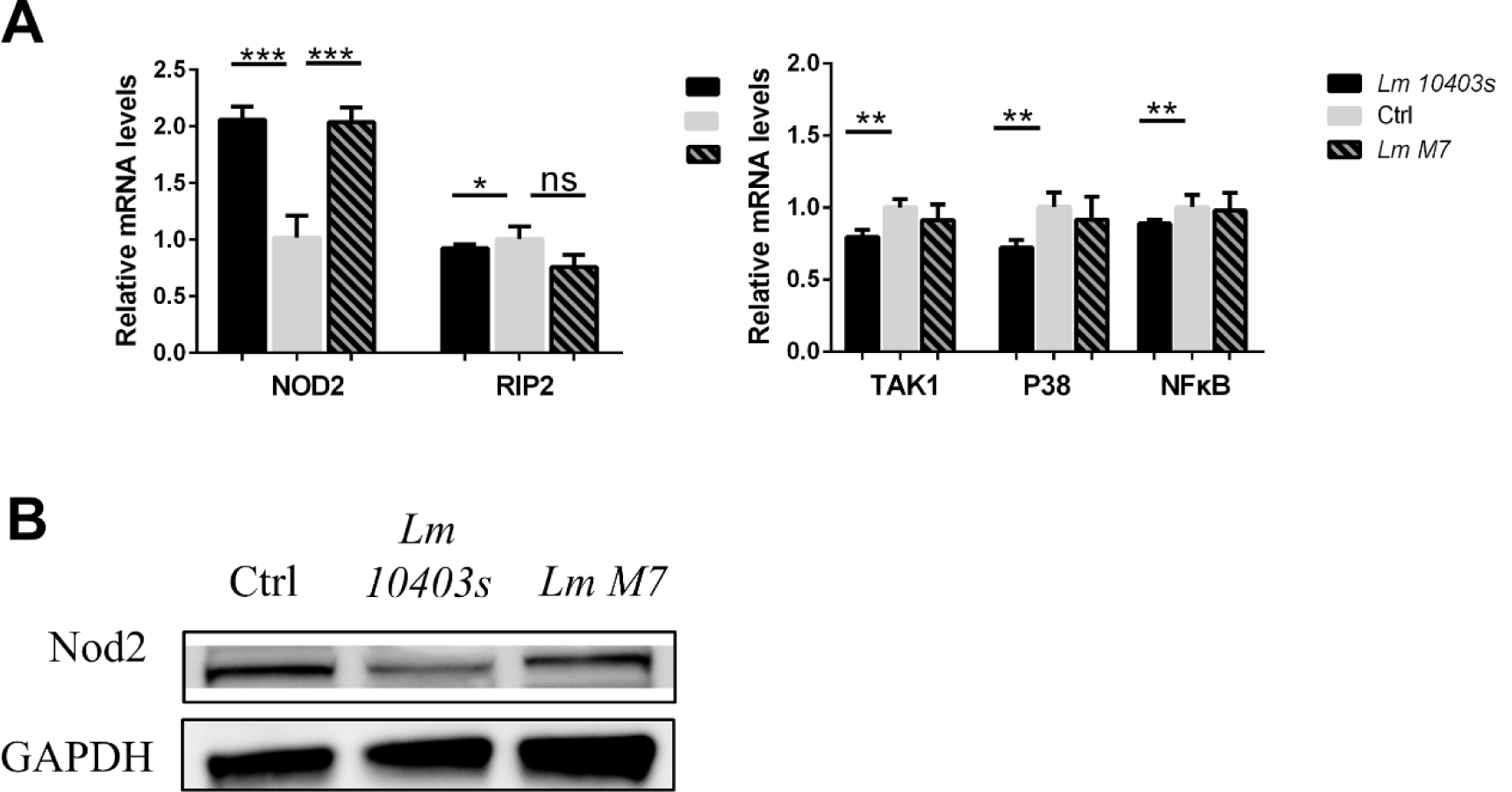
Expression levels of the DEPs were verified using RT-PCR and western blot analysis. A. Verification results of Nod2 pathway genes (Nod2, RIP2, TAK1, P38, and NF-κB) at the mRNA levels. B. Verification results of Nod2 at the protein levels.

## Discussion

Foodborne illness has been a major threat to human health and public health ^[48]^. Although the incidence has been greatly reduced with the improvement of sanitary conditions, food poisoning is still a major problem today ^[49]^. Listeriosis is one of the most serious foodborne diseases, and mainly caused by contaminated food. *Listeria monocytogenes* enters the digestive tract through contaminated food, and then crosses the intestinal barrier, which is a critical step in systemic infection ^[26]^. In the researches on the stage of intestinal infection, most analyzed virulence gene and survival mechanism of *L. monocytogenes*, and the host response ^[50, 51]^. Furthermore, the infection may alter host intestinal microbiota to promote bacterial colonization ^[25]^. In the proteomics analysis of *L. monocytogenes*, many studies focused on bacterial proteins, and explored the relationship between some proteins and bacterial virulence by comparing the different protein expression between different strains in stress, biological metabolism, and virulence genes ^[23, 39]^. The others focused on host protein changes, and investigated the interaction between host response and bacterial virulence. _^[52]^_.

However, researches on changes of intestinal epithelial host protein were not complete. Previous studies used either animal models or single intestinal cell line, which did not exclude the effects of immune cells or could not fully represent epithelial cells. As an emerging intestinal model, small intestine organoids have been used to study the interaction between bacteria and hosts ^[47, 53]^. However, in the study of *L. monocytogenes* and small intestine organoids, the focus was only to verify whether the intestine organoids could be used as an invasion model for *L. monocytogenes* and the apparent damage effect of bacteria on organoids. On the other hand, there was no comprehensive analysis of infected intestine organoids using proteomics.

In the present study, tandem mass tag-based quantitative proteomic analysis was used to compare the total proteomes in organoids infected by different toxic *L. monocytogenes*. Quantitative analysis demonstrated 154 differentially expressed proteins in the virulent strain (*L. monocytogenes 10403s)*-infected organoids, 213 proteins in the low virulent strain (*L. monocytogenes M7)*-infected organoids, and 62 proteins in the *L. monocytogenes 10403s* vs *L. monocytogenes M7* comparison group. These proteins were found to be involved in cell transport, binding, biological metabolism, energy metabolism, transcriptional regulation, signal transduction, and defense response.

The infection of *L. monocytogenes* in intestinal organoids is a process that involves many proteins and pathways. The results of analyzing *L. monocytogenes 10403s* and *L. monocytogenes M7* groups showed that the damage of *L. monocytogenes* infection was mainly reflected in the destruction of intestinal barrier, affecting the disease-signaling pathway, and changing the metabolic process of host cell. In addition, different toxic *L. monocytogenes* led to the changes of five important pathways related to host immune process.

ECM-receptor interaction is a micro-environmental pathway that maintains cell and tissue structure and function, which leads to a direct or indirect control of cellular activities such as adhesion, migration, differentiation, proliferation, and apoptosis. Recent studies have identified that this pathway was possibly involved in the development of breast cancer ^[54]^. Studies confirmed that these proteins were utilized by pathogens to adhere to and invade host tissues, and can increase adherence of *L. monocytogenes* to HEp-2 cells ^[55, 56]^. *L. monocytogenes* upregulated many ECMs during the infection, such as fibronectin, Laminin, and collagen, indicating that *L. monocytogenes* can improve the ability of adhesion and invasion of cells by adjusting ECM.

The significantly different proteins in HIF-1 signaling pathway and Ferroptosis were Tfrc and Tf, which regulated intracellular iron. They were both upregulated in *L. monocytogenes 10403s* and *L. monocytogenes M7* infection groups. It was found that transferrin (Tf), transferrin receptor (Tfrc) and ferroportin favored oxidative damage and Ferroptosis by increasing iron uptake and reducing iron export ^[57]^. However, hypoxia inducible factor-1 (HIF-1) was not identified, which may be caused by the lack of immune cells in the small intestine organoids. Therefore, it indicated that the enriched HIF-1 signaling pathway was not caused by HIF-1. In addition, there was no significant difference in the key regulatory protein glutathione peroxidase 4 (Gpx4) in the Ferroptosis pathway. This indicated that the effect of different toxic *L. monocytogenes* on Ferroptosis was not critical.

Daniel G. et al. claimed that complement was an essential defense of *L. monocytogenes* infection ^[58]^. Complement, coagulation and fibrinolytic systems can form serine protease system, which plays an essential role in the innate immune responses. The interplay between complement and coagulation contributed to strengthen innate immunity, and activate adaptive immunity to eliminate bacteria ^[59]^. In our study, three fibrinogen chains (Fga, Fgb and Fgg) were upregulated in two *L. monocytogenes* infections, while coagulation factor X (F10) and C3 was upregulated only in *L. monocytogenes M7* infection group. It indicated that the different toxic *L. monocytogenes* activated the complement system and coagulation cascade in the stage of intestinal infection, and low-virulence strains caused a more significant coagulation cascade.

The NOD-like receptor signaling pathway is mediated by Nod-like receptors in the host cell and is also an important innate immune response ^[60]^. Nod2 in NOD-like receptors can be expressed in intestinal epithelial cells, such as Paneth cells and stem cells ^[61]^. Besides, extensive studies have shown that Nod2 plays an important role in maintaining the balance between bacteria, epithelial cells and the innate immune response of the host ^[31]^. Nod2 recruits downstream proteins by recognizing the muramyl dipeptide (MDP) in the cell wall of pathogens, and then induces the activation of NF-κB, MAPK, and caspase-1 pathways^[62]^. In general, bacteria will activate Nod2 during infection and increase the expression level of Nod2. Interestingly, Nod2 was down-regulated in both *L. monocytogenes* 10403 and *L. monocytogenes* M7 groups in our result.

Studies showed that under the stimulation of different concentrations of MDP, Nod2 in dental pulp stem cells was activated, but the expression level was reduced ^[63]^. This suggested that Nod2 in stem cells could be inhibited by MDP. In addition, studies showed that the expression of Nod2 in the terminal ileum of sterile mice was lower, and the expression of Nod2 was increased after supplementing symbiotic bacteria in sterile mice ^[64]^. This indicated that the expression level of Nod2 in the intestine was related to commensal bacteria. Therefore, the reason for the low expression of Nod2 in the small intestine organoid model could be related to the lack of symbiotic bacteria.

Combined with the mRNA expression results of Nod2 pathway proteins, it can be seen that the infection led to the activation of Nod2 at the mRNA level, while it caused a reduction in the protein expression. This indicated that the bactericidal effect of Nod2 pathway in intestine organoids was limited within 18h. It could not effectively eliminate *L. monocytogenes*, and resulted in a significant reduction of Nod2 expression in damaged cells (such as intestinal stem cells). Therefore, the overall protein level of Nod2 and mRNA expression of other proteins in the Nod2 pathway were decreased in *L. monocytogenes 10403s* group.

Besides, NOD2 is highly expressed in Paneth cells, which defense intestinal pathogen by secreting antimicrobial compounds. Several studies highlighted the essential role that NOD2 played in maintaining the equilibrium between intestinal microbiome and host immune responses ^[61, 65]^. In addition, recent studies showed that Nod had an important effect on intestinal stem cells. Nigro et al. reported that NOD2 provided cytoprotection to intestinal stem cells, and Levy et al. found that the mechanism of NOD2-mediated cytoprotection involved the clearance of the lethal excess of ROS molecules through mitophagy ^[66, 67]^. Combined with our unpublished research results, it was found that the damage of *L. monocytogenes* to intestinal stem cells could be achieved by down-regulation of NOD2.

Moreover, the downstream of the NOD-like receptor-signaling pathway includes the formation of inflammatory corpuscle complexes and the activation of caspase-1. Pycard (also known as ASC) is a key adaptor protein of inflammatory bodies (such as NLRP3) and an essential protein that activates inflammatory responses and apoptosis signaling pathways ^[68]^. In the *L. monocytogenes 10403*s and *L. monocytogenes M7* groups, Pycard expression was down-regulated, indicating that *L. monocytogenes* may down-regulate Pycard to inhibit the formation of inflammatory corpuscle complexes. In addition, in the *L. monocytogenes 10403* vs *M7* comparison group, Txn2 and Nek7 related to NLRP3 inflammatory body assembly were significantly down-regulated. Specifically, they were down-regulated by *L. monocytogene*s 10403s and up-regulated by *L. monocytogenes M7*, while their change difference did not reach 1.2 times significantly. This showed that that difference between the two strains on the Nod-like receptor-signaling pathway in the host was mainly in the regulation of inflammasomes assembly.

Overall, KEGG enrichment analysis showed that different toxic *L. monocytogenes* increased the expression of adhesion and invasion-related proteins, reduced energy metabolism of host and triggered various host defense responses. Besides, the virulent strain *L. monocytogenes 10403*s had a more significant activation effect on the Ferroptosis pathway, while the low virulent strain *L. monocytogenes M7* has a more significant activation effect on the complement system. More importantly, *L. monocytogenes* with different toxicity could affect the proliferation and cell protection of intestinal stem cells by down-regulating Nod2. In addition, the down-regulated proteins of *L. monocytogenes 10403s* vs *L. monocytogenes M7* comparison group were enriched into systemic lupus erythematosus and transcriptional misregulation in cancer. It suggested that the low virulent strain had a stronger interference effect on immunodeficiency disease and transcriptional regulation.

In summary, complex responses to virulent and low virulent *L. monocytogenes* infections were revealed by TMT-based quantitative proteomics analysis using intestinal organoids. The DEPs between the *L. monocytogenes 10403s* and *L. monocytogenes M7* infected groups displayed similar biological functions and subcellular localizations as previous analysis. The difference in their influence on the host biological function was mainly reflected in transcription regulation and metabolism. These different DEPs were mainly distributed in the nucleus, and their domains were related to histones. Furthermore, complement and coagulation cascade and NOD-like receptor-signaling pathway were detected as the innate immune responses caused by two strains. Our result revealed that the modulation of protein expression attributed to the strategy of *L. monocytogenes* to overcome host defense response, and the data may give a comprehensive resource for investigating the overall response of intestinal epithelial cells excluding immune cells to infection with different toxic *L. monocytogenes*.

## Acknowledgements

The authors would like to thank Prof. Weihuan Fang for the *Listeria monocytogenes* strain, and Prof. Qinghua Yu for giving suggestions to the cultured system of mouse small intestinal organoids. This work was supported by the Fundamental Research Funds for the Central Universities (KYZ201823 and KYYZ201803) and Jiangsu Agriculture Science and Technology Innovation Fund (CX(18)2024).

---

**Table S1.**
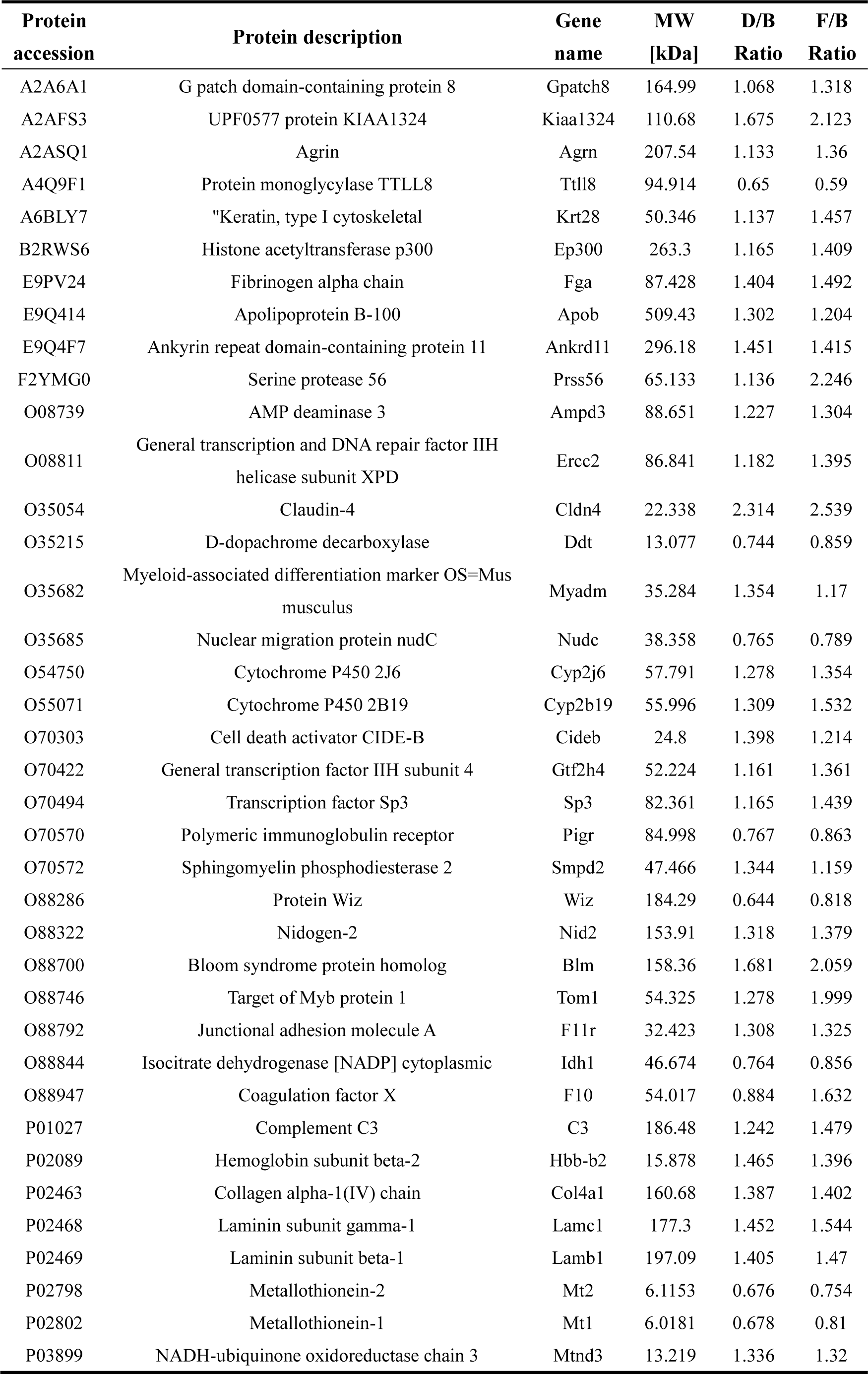
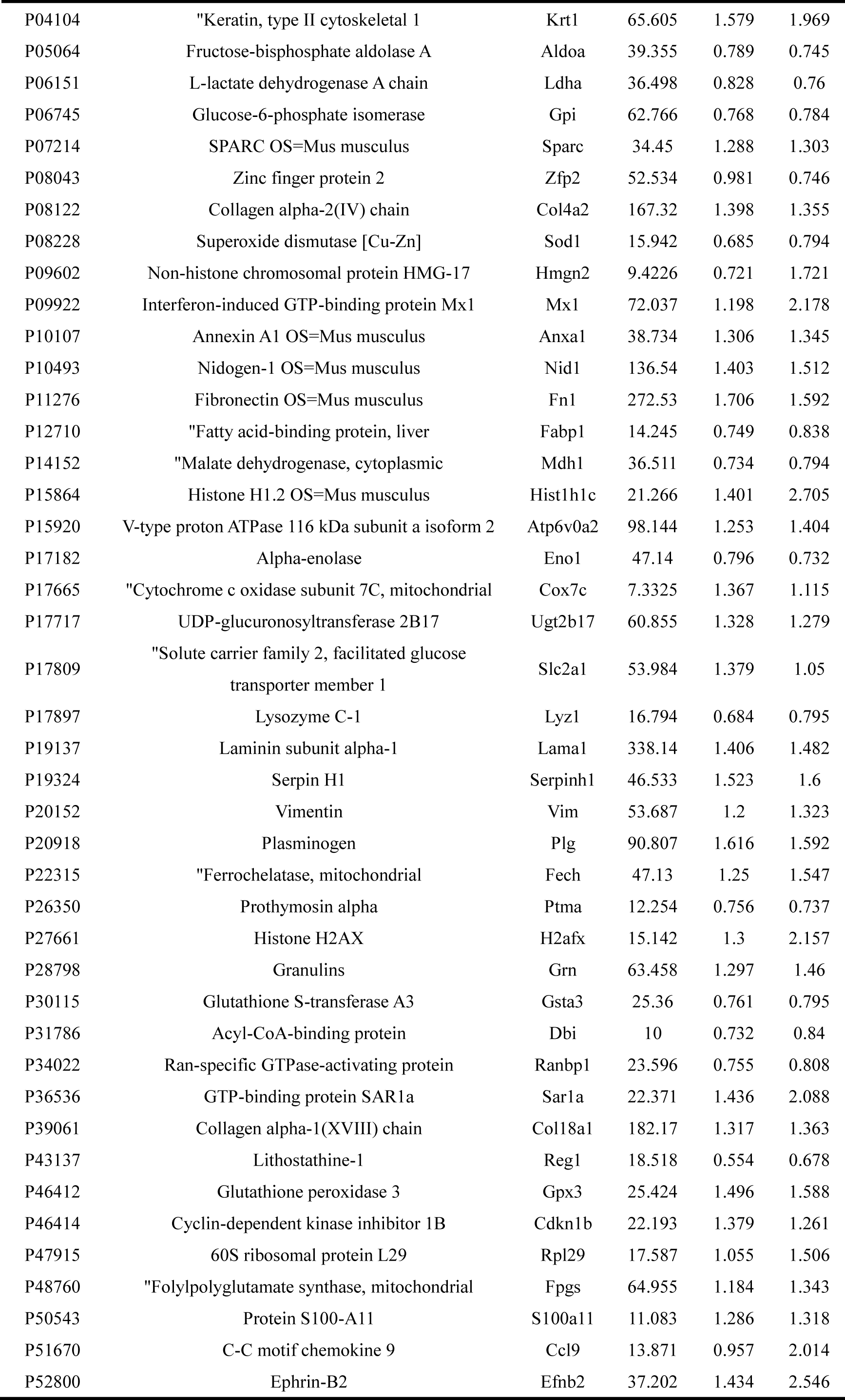
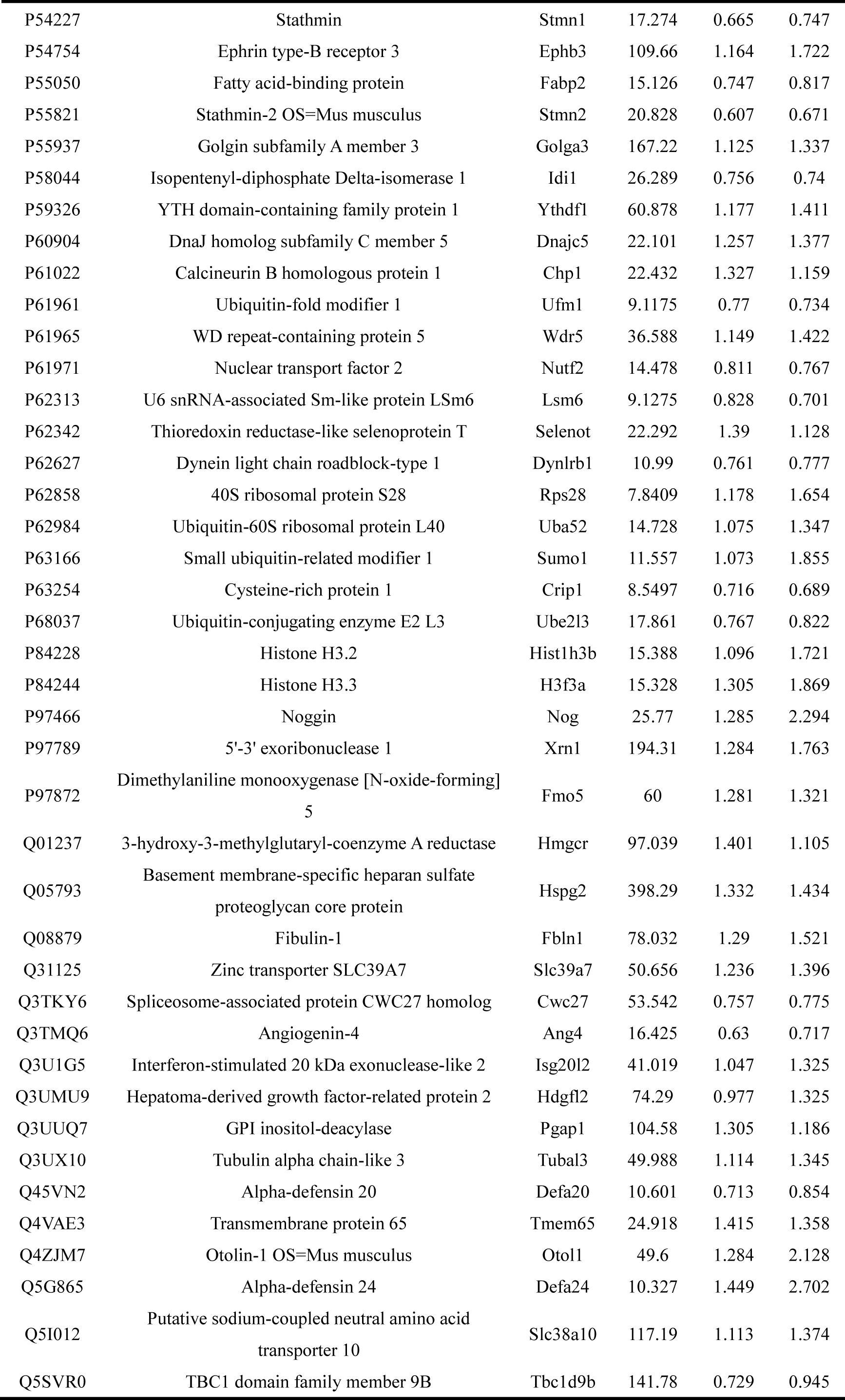
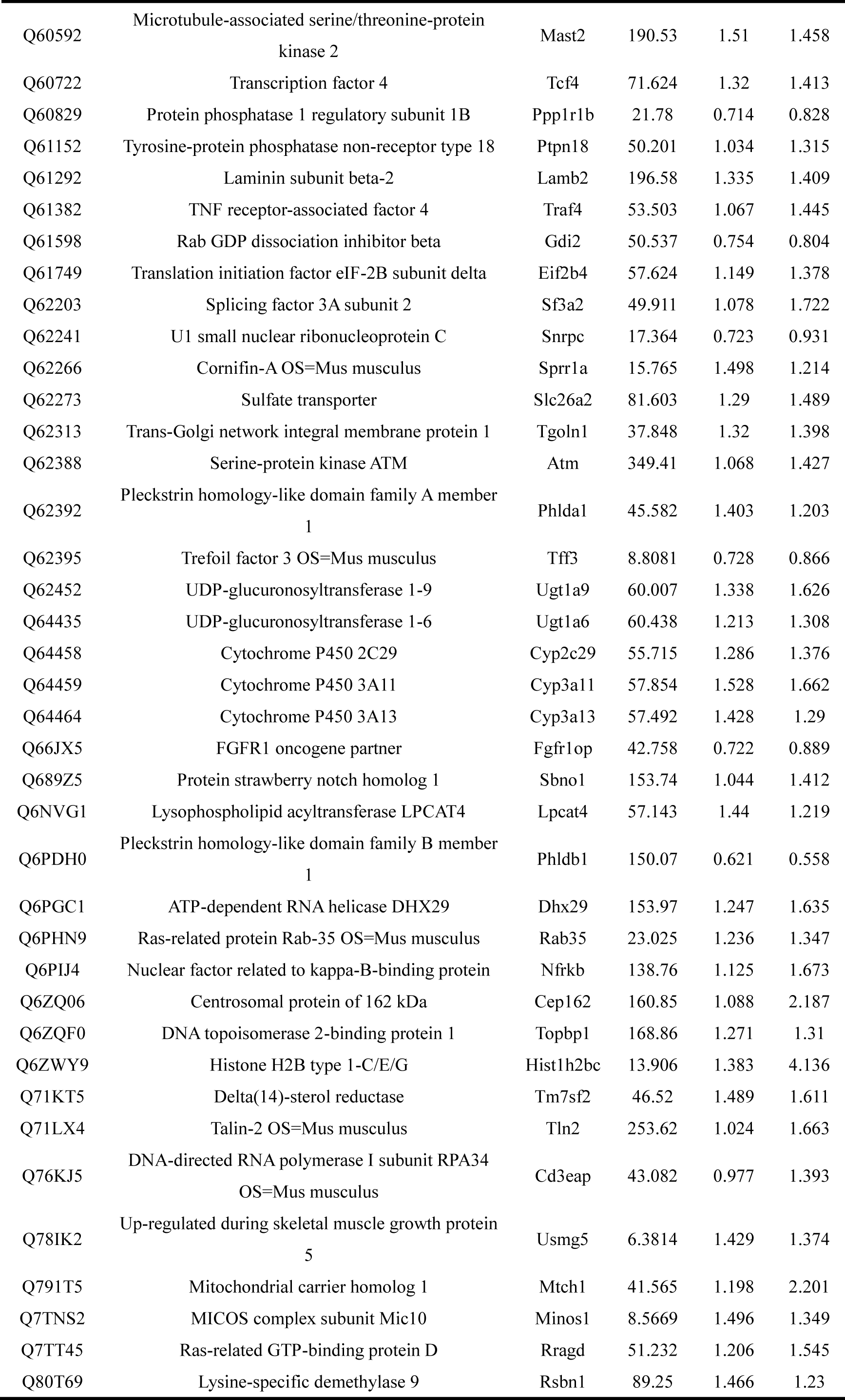
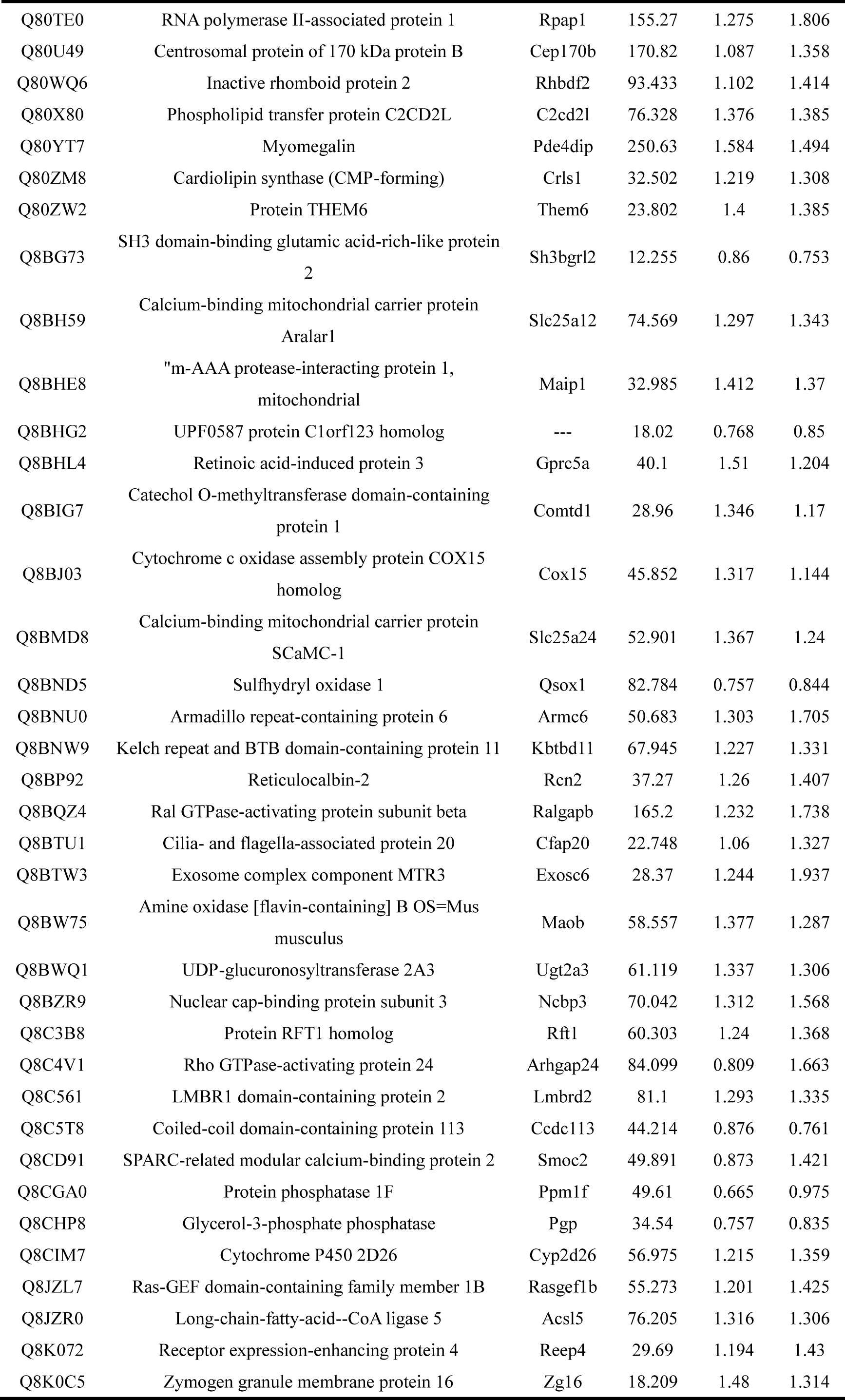
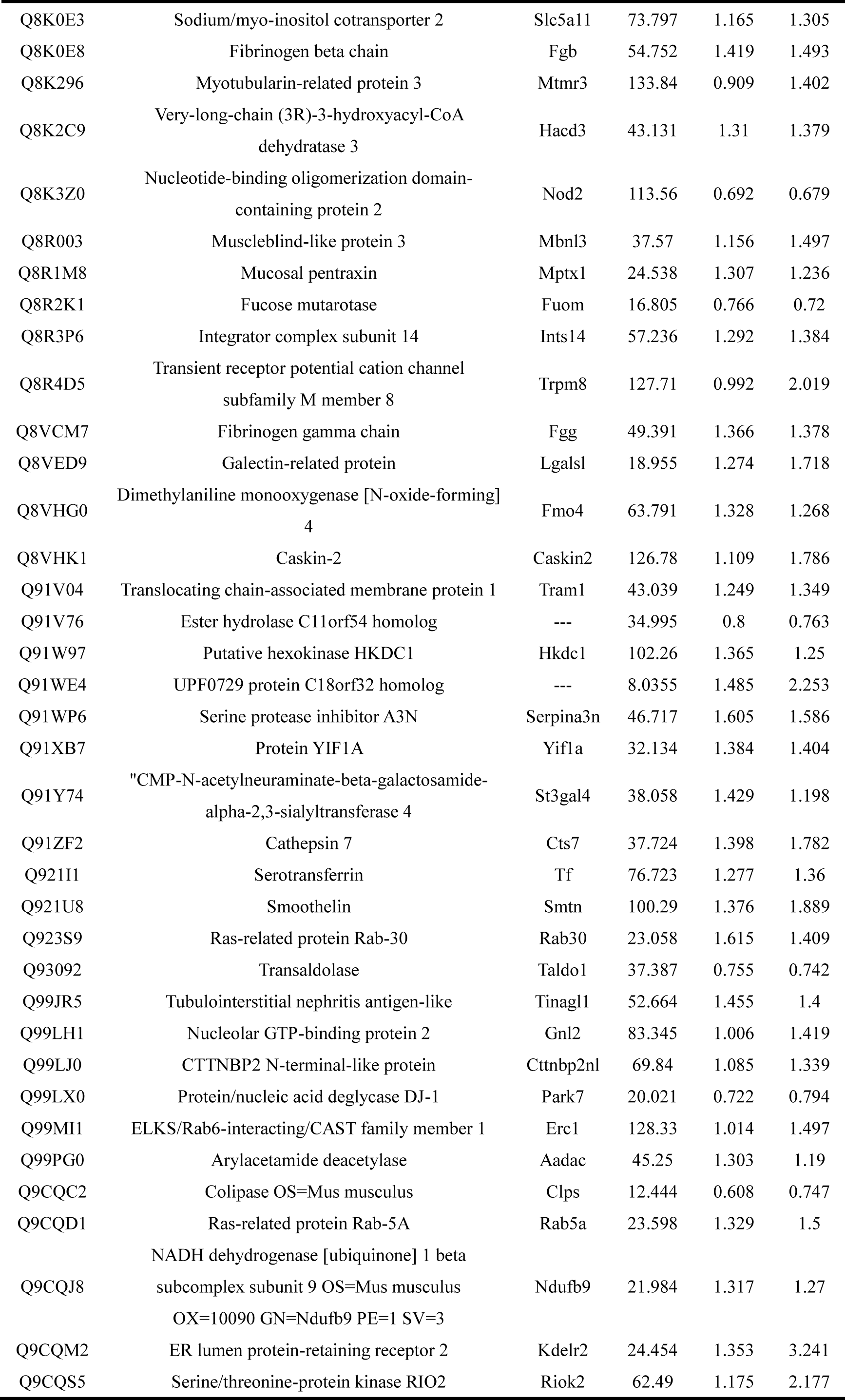
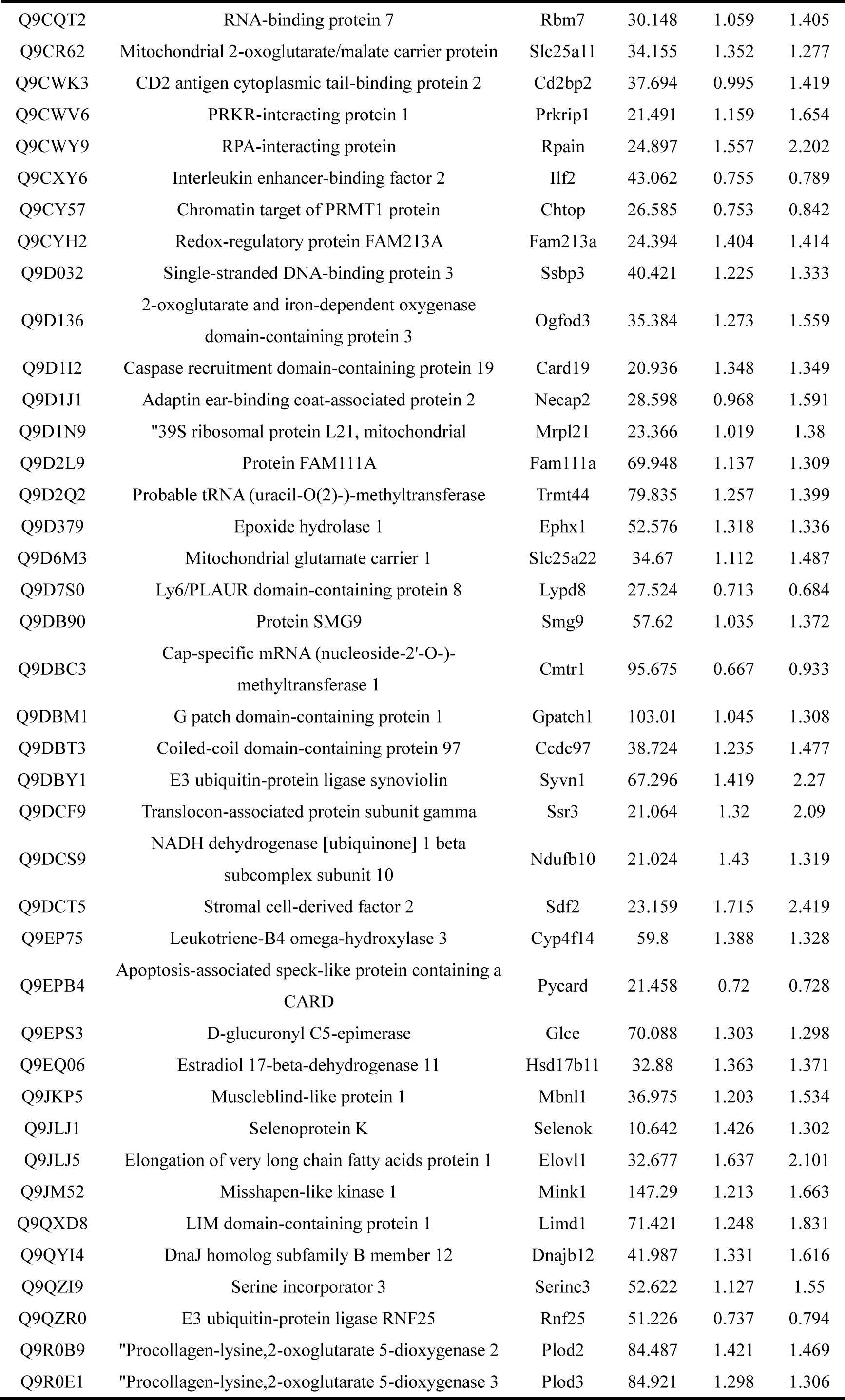
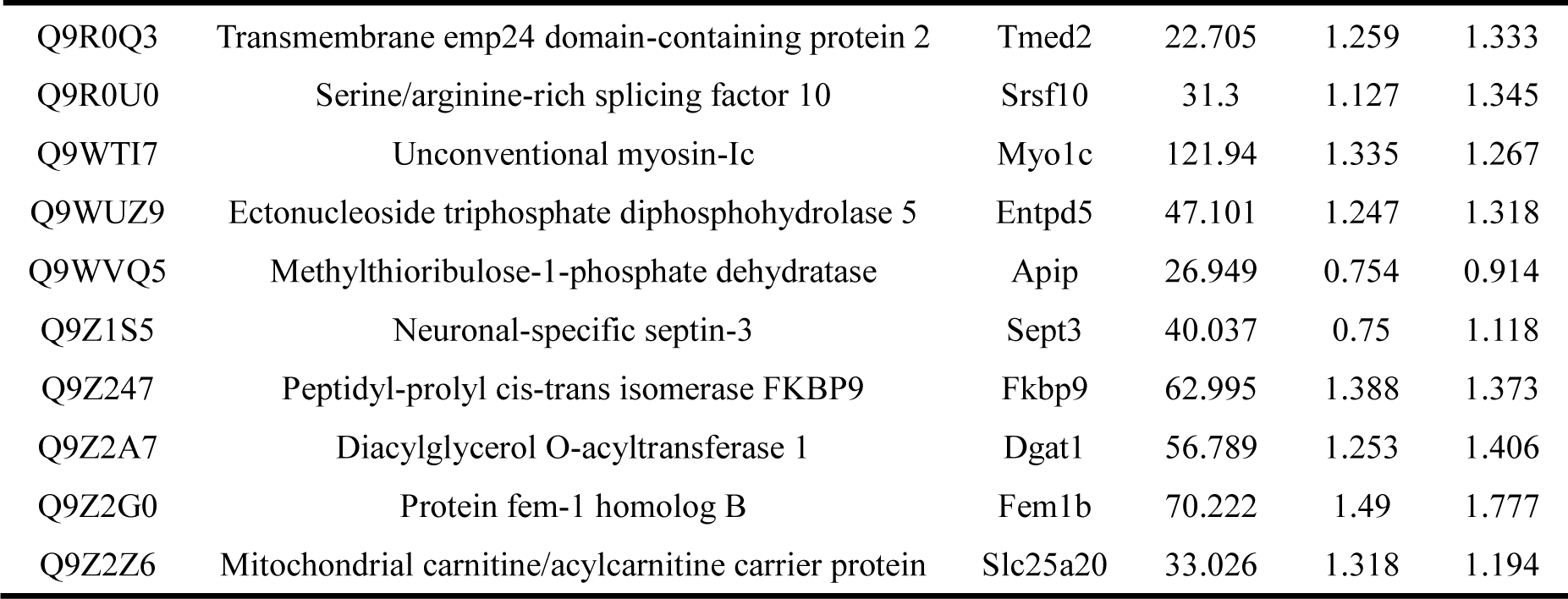
Relative expression of DEGs in *Lm 10403s* vs Control (D/B) and *Lm M7* vs Control (F/B)

